# Neuroanatomical substrates in Parkinson’s Disease psychosis and their association with serotonergic receptor gene expression: A coordinate-based meta-regression analysis

**DOI:** 10.1101/2022.11.14.516465

**Authors:** Sara Pisani, Brandon Gunasekera, Yining Lu, Miriam Vignando, Dominic ffytche, Dag Aarsland, K. Ray Chaudhuri, Clive Ballard, Jee-Young Lee, Yu Kyeong Kim, Latha Velayudhan, Sagnik Bhattacharyya

## Abstract

**Background:** Common neural underpinning of Parkinson’s Disease (PD) psychosis across different structural magnetic resonance imaging (MRI) studies remains unclear to this day with few studies and even fewer meta-analyses available.

**Objectives:** Our meta-analysis aimed to identify and summarise studies using MRI approach to identify PD psychosis-specific brain regions and examine the relation between cortical volume loss and dopaminergic and serotonergic receptor density.

**Methods:** PubMed, Web of Science and Embase were searched for MRI studies of PD psychosis (PDP) compared to PD patients without psychosis (PDnP). Seed-based *d* Mapping with Permutation of Subject Images was applied in the meta-analysis where coordinates were available. Multiple linear regressions to examine the relationship between grey matter volume loss in PDP and receptor gene expression density (extracted from the Allen Human Brain Atlas) were conducted in R.

**Results:** We observed lower grey matter volume in parietal-temporo-occipital regions from our meta-analysis (N studies =10, PDP n=211, PDnP, n=298). These results remained significant after adjusting for PD medications and for cognitive scores. Grey matter volume loss in PDP was associated with local expression of 5-HT1a (b=0.109, *p*=0.012) and 5-HT2a receptors (b=-0.106, *p*=0.002) also after adjusting for PD medications (5-HT1a, *p* = 0.005; 5-HT2a, *p* = 0.001).

**Conclusions:** Widespread cortical volume loss in the parieto-temporo-occipital regions involved in information processing and integration, as well as attention, could result in PD psychosis symptoms. Neurobiological mechanisms implicating serotonergic receptors may also contribute to this condition.

## 1. Introduction

Non-motor symptoms of Parkinson’s Disease (PD) are distressing, debilitating, and associated with poor quality of life in PD patients. Specifically, hallucinations and delusions, the key manifestations of PD psychosis (PDP), negatively impact patients and caregivers and increase care burden and risk of hospitalisation (Aarsland et al., 2007; Martinez-Martin et al., 2015). Although initially considered a by-product of dopamine augmentation treatments, evidence suggests that drug-naïve PD patients may also experience psychosis symptoms (Friedman, 2016; Pagonabarraga et al., 2016). Severity and duration of PD, dopaminergic medications, sleep disorders, cognitive decline, widespread Lewy Body pathology, and late PD onset can be risk factors for developing psychosis symptoms (Barrett et al., 2018; Chang & Fox, 2016; Factor et al., 2014; Ffytche et al., 2017; Williams & Lees, 2005).

The mechanisms underlying the emergence of psychotic symptoms in PD remain unclear. Results from structural as well as task-based and resting state magnetic resonance imaging (MRI) and positron emission tomography (PET) studies have generally reported grey matter reductions (Bejr-Kasem et al., 2019; Gama et al., 2014; Nishio et al., 2018; Ramírez-Ruiz et al., 2007) encompassing the ventral and dorsal visual pathways and hippocampus (Alzahrani & Venneri, 2015; Lenka et al., 2015; Yao et al., 2016) as well as altered temporo-parieto-occipital activation, metabolism or functional and structural connectivity in large-scale brain networks (Boecker et al., 2007; Hall et al., 2019; Hepp et al., 2017; Lefebvre et al., 2016; Shine, Halliday, et al., 2014; Shine et al., 2015; Stebbins et al., 2004). This is consistent with a recent meta-analytic synthesis reporting fronto-temporo-parieto-occipital brain grey matter reduction in a mixed group of participants with PD and Lewy body dementia (DLB) with visual hallucinations (Pezzoli et al., 2021), although they did not consider the effect of potential confounders such as PD medication or concomitant symptoms, e.g., cognitive decline. Complementing this, a recent mega-analysis (Vignando et al., 2021) has reported extensive reduction of whole-brain cortical thickness and surface areas in visual cortex, left insula and hippocampus in PD patients with visual hallucinations compared to those without, as well as a relationship between cortical thickness loss and higher regional availability serotonergic (5-HT2a and 5-HT1a) and dopaminergic receptors (D2/D3). Collectively, these reviews focused on related different metrics of brain structure (volume, cortical thickness and surface). Meta-analyses integrating evidence regarding alterations in brain areas in PD psychosis may help unravel neuroanatomical substrates underpinning Parkinson’s psychosis. Here, we have conducted a quantitative analysis of structural neuroimaging studies investigating the neural correlates of psychosis in PD. Further, we examined whether the spatial distribution of brain structure alteration was associated with the spatial architecture of brain expression of the genes for dopaminergic (D1 and D2) and serotonergic receptors (namely 5-HT2a and 5-HT1a), the key candidate pathways implicated in PD psychosis (Hacksell et al., 2014). Unlike the approach adopted by Vignando et al. (2021) where they examined the relationship between the structural neuroimaging correlate and the density distribution data for each receptor of interest separately, we included the key candidate receptors of interest in a multiple regression model to allow investigation of their relationship with neuroimaging correlates after taking into account the effect of receptors belonging to other candidate pathways of interest.

## 2. Methods

### 2.1. Search strategy and eligibility criteria

We followed (PROSPERO registration number: CRD42020221904) the Preferred Reporting Items for Systematic Reviews and Meta-Analyses (PRISMA) guidelines (Moher et al., 2009) and a detailed description of the search strategy is outlined in Supplementary Material 1. Studies identified from PubMed, Web of Science, Embase and the Neurosynth database were included if they examined brain alterations associated with psychosis symptoms (i.e., hallucinations and/or delusions) after PD diagnosis using different neuroimaging modalities, and a case-control design and provided brain coordinates (or available statistical maps) in standardised reference spaces, e.g., Montreal Neurological Institute (MNI) or Talairach space. There were no exclusion or restriction for this search. For this meta-analysis, we solely focused on studies that used structural magnetic resonance imaging (MRI) modality.

### 2.2. Data extraction

Data extraction involved: study details (i.e., authors’ names, year of publications), study design, scanner characteristics (e.g., manufacturer), sample size, sample characteristics (e.g., age, gender, education levels), PD onset age, disease duration, PD symptoms, clinical measures of psychosis symptoms, cognitive measures, dopamine-replacement medications (expressed in Levodopa equivalent daily dose (LEDD)), and other clinical outcome measures (e.g., depression). Where only median values were available, this was converted into mean (Luo et al., 2018). Coordinates type (e.g., MNI or Talairach space), and associated *t* values were also extracted. Where coordinates were expressed in Talairach or other standard normalised spaces, they were converted into MNI coordinates. Authors were contacted for studies that did not report coordinate data. Data extraction was conducted independently by two researchers (SP, YL). Discrepancies were addressed through consensus or discussion with senior researchers.

### 2.3. Data synthesis

A coordinate-based meta-analysis was conducted for structural MRI studies reporting voxel-based morphometry (VBM) results using peak coordinates and/or statistical maps by employing a random-effects approach using Seed-based *d* Mapping with Permutating Subject Images (SDM-PSI) (version 6.21, https://www.sdmproject.com/) (Albajes-Eizagirre et al., 2019). This involved computation of upper and lower bounds of effect size maps for studies based on the available peak coordinates by imputing voxel-wise effect size (Albajes-Eizagirre et al., 2018) and combining them with statistical parametric maps provided by the studies (where applicable) converted into effect size maps to carry out a random-effects meta-analysis. Contrasts of interest were ‘PD psychosis (PDP) < PD without psychosis (PDnP)’ (i.e., grey matter loss) and ‘PD psychosis (PDP) > PD without psychosis (PDnP)’ (i.e., increased grey matter) with and without controlling for the effect of PD medications (expressed in LEDD), and cognitive scores. Due to the various neurocognitive scales used to assess cognitive abilities in PDP and PDnP patients, we computed standardised scores for each study by dividing the mean score by the standard deviation (SD) for each patients’ group. Statistical parametric maps provided by the studies (where applicable) were registered to the SDM template, and their *t* values were converted into effect sizes and included in the analysis. SDM-PSI provides *I^2^* statistic to assess heterogeneity in the observed peaks as well as between-study heterogeneity (Higgins et al., 2003). Funnel plots for each peak and Egger’s test (Egger et al., 1997) were used to assess publication bias. Clusters smaller than 10 voxels were discarded.

We examined the association between grey matter volume loss (measured using Hedges’ *g* effect-size estimates extracted from the centroid of the SDM-PSI meta-analytic map parcellated across 78 brain regions of the Desikan-Killiany atlas (Desikan et al., 2006)) in PDP compared to PDnP patients with dopaminergic (D1/D2) and serotonergic (5-HT2a/5-HT1a) mRNA microarray gene expression using a multiple linear regression analysis. A secondary analysis included LEDD as a covariate when computing Hedges’ g effect-size estimate. Gene expression data for dopaminergic and serotonergic receptors was extracted from the Allen Human Brain Atlas which includes microarray expression data in tissue samples from six healthy adult human brains with more than 20,000 genes quantified across cortical and subcortical regions, including brainstem and cerebellum (Hawrylycz et al., 2012). Probe-to-gene re-annotation data was downloaded using the information from Arnatkeviĕiūtė et al. (2019). Sample-to-region matching, and extraction of the gene expression data followed established approaches (Arnatkeviĕiūtė et al., 2019; Markello et al., 2021). Gene expression data for D3 receptor could not be matched to the probe and therefore was not extracted.

### 2.4. Quality rating assessment

Study quality was assessed with the Newcastle-Ottawa rating Scale (Wells et al., 2000) for case-control studies. This scale includes three methodological domains: “Selection” (i.e., definitions of cases and controls, and their selection), “Comparability” (i.e., comparison of cases and controls on the variables of interest, inclusion of covariates), and “Exposure” (i.e., how exposure such as condition was defined and ascertained). This rating is based on a star system: studies can be awarded a maximum of one star for each “Selection” and “Exposure” items, and a maximum of two stars for the “Comparability” domain. Quality ratings were conducted by one researcher. In this review, neural substrates of PDP patients were compared to those in PDnP patients, thus the latter acted as control or comparison group. Therefore, the items “Selection of controls” and “Definition of controls” were specifically related to PDnP patients.

## 3. Results

### 3.1. Study characteristics

Our systematic search identified 14 studies out of which 9 reported peak coordinate data, and one provided T maps (Bejr-Kasem et al., 2019; Bejr-Kasem et al., 2020; Firbank et al., 2018; Goldman et al., 2014; Lawn, 2021; Lee et al., 2017; Pagonabarraga et al., 2014; Ramírez-Ruiz et al., 2007; Shin et al., 2012; Watanabe et al., 2013). Thus, we included a total of 10 published articles (Fig. 1) reporting on a total of n= 211 PDP (mean age ± SD = 69.01 ± 4.90; motor symptom score, mean ± SD = 28.72 ± 11.69, LEDD mg/day mean ± SD = 651.9 ± 318.14) and n=298 PDnP patients (mean age ± SD = 67.34 ± 4.59, motor symptom scores, mean ± SD = 25.94 ± 8.63, LEDD mg/day mean ± SD = 577.1 ± 263.11) (Table 1).

**Figure 1.**
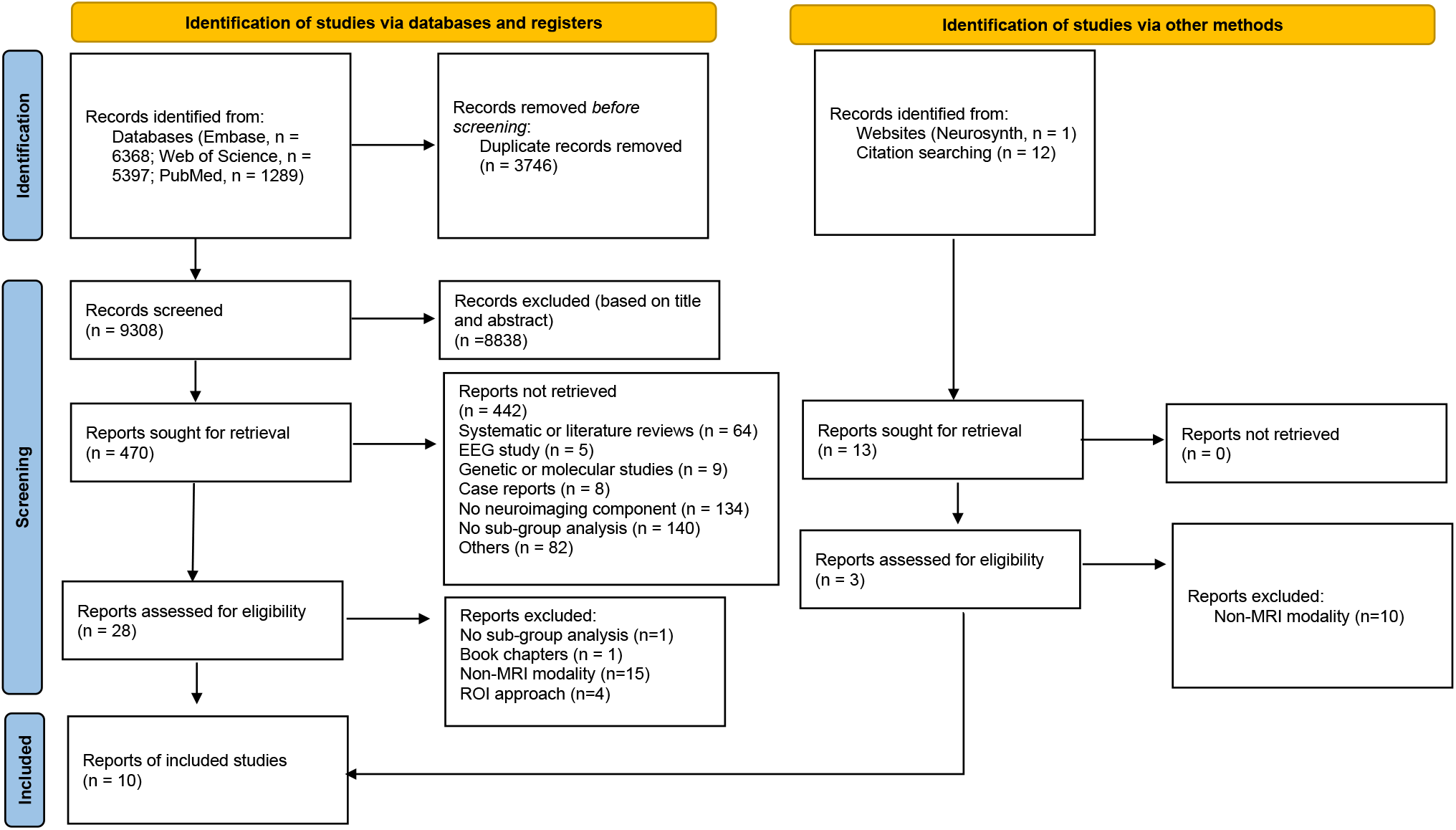
PRISMA flowchart showing study selection procedures.

**Table 1.**
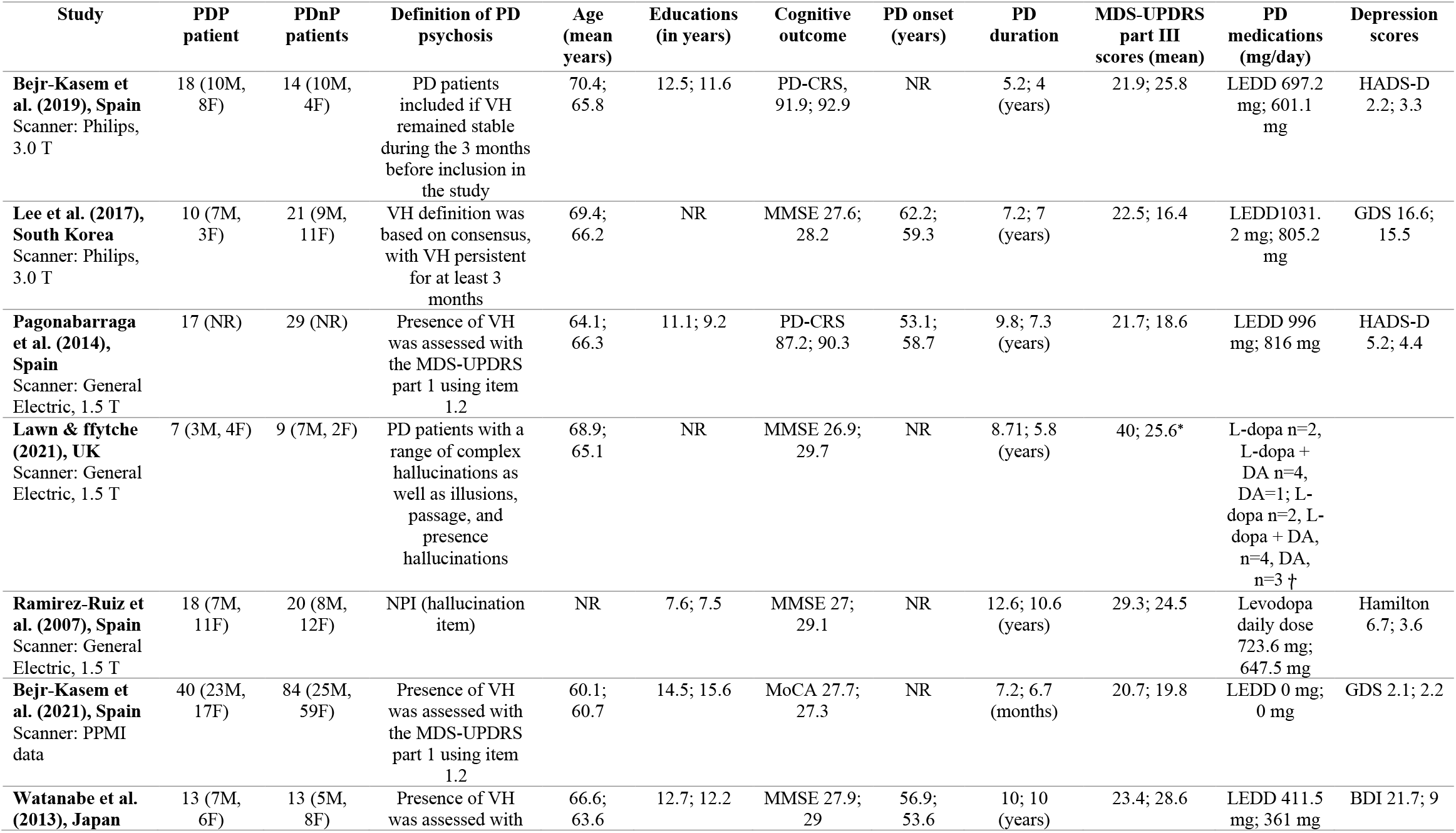
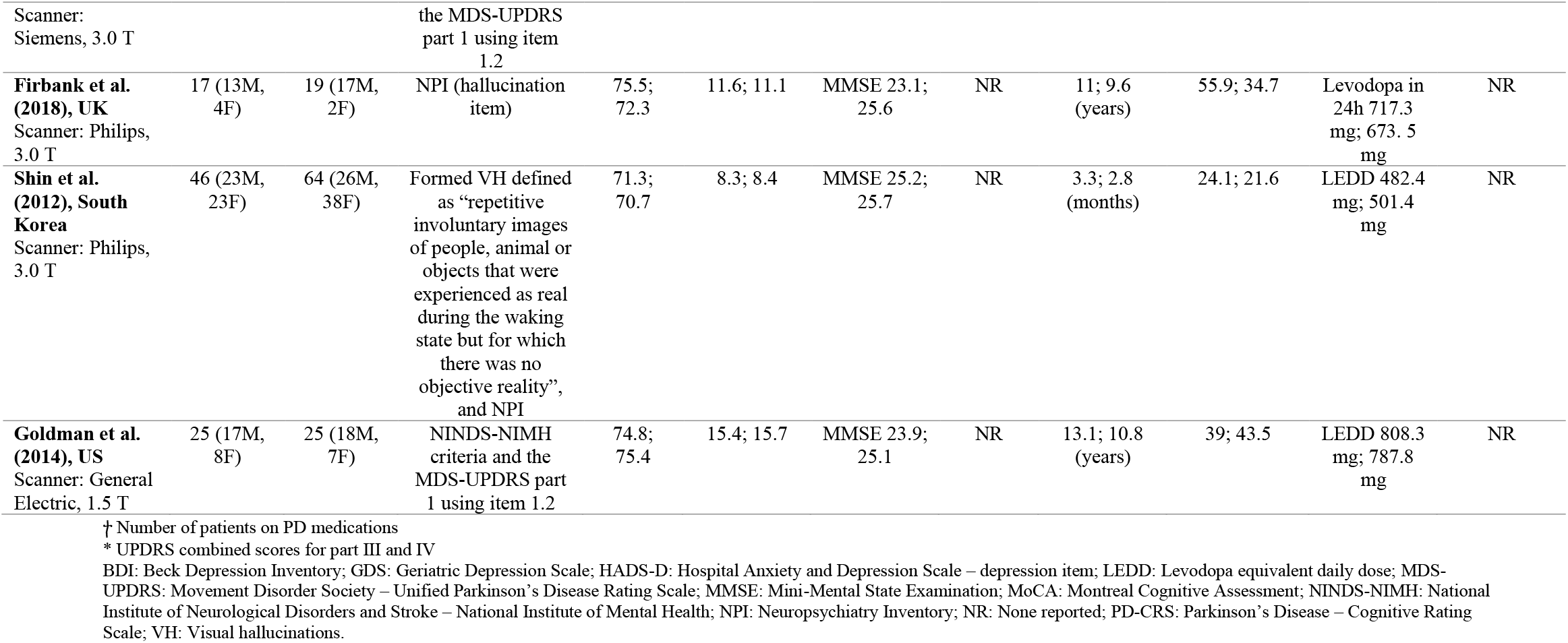
Study description with statistics (mean, unless otherwise specified) presented for each patient group (PDP patients; PDnP patients). Number of patients for each group are presented alongside gender (males, M; females, F), years of education, scores on cognitive outcomes. Clinical variables of interest are also reported, e.g., motor symptoms (according to the MDS-UPDRS), PD medications (expressed in mean daily mg where available), and depression scores where available.

Full quality ratings are reported in Supplementary Material 2 (eTable1). Briefly, all studies had full score on the “Exposure” domain (i.e., ascertainment, non-response rate and ascertainment method for PDP and PDnP patients); “Comparability” was good in all studies due to their matched design, whereby patients were matched on age, gender and other clinical or demographic variables, whilst other studies reported these variables as covariates in the analysis. “Selection” domain included selection and definition of both PDP patients (i.e., cases) and PDnP patients (i.e., controls), selection of PDP patients was assigned one star (i.e., the maximum) in all studies. Similarly, selection of PDnP patients was assigned one star in all studies, whilst definition of such group was clear in one study which was assigned one star in this domain.

### 3.2. Meta-analysis: Voxel-based morphometry (VBM)

Meta-analysis of 10 MRI studies (one provided statistical maps (Lee et al., 2017)) showed that PDP patients had reduced grey matter volume in parietal-temporo-occipital areas compared to PDnP patients with the largest clusters located in the right precuneus (extending to left precuneus and bilateral cuneus; voxel number = 1059, Z=-2.990, *p* = 0.001), bilateral inferior parietal gyrus (left inferior parietal gyrus, extending to left angular gyrus and supramarginal gyrus; voxel number = 296, Z=-2.449, *p* = 0.007; right inferior parietal gyrus, extending to right angular gyrus; voxel number = 277, Z=-2.225, *p* = 0.013), left inferior occipital gyrus (voxel number = 262, Z =-2.627, *p* =0.004), and right middle temporal gyrus (extending to right inferior temporal gyrus; voxel number = 225, Z = −2.664, *p* = 0.003) (eFigure1, Supplementary Material 3). We did not observe any significant peaks in the opposite direction (i.e., ‘PDP > PDnP’). Neither between-study heterogeneity as a proportion of total variability (all *I^2^* statistic <25%, except right inferior parietal gyrus, which was ~27.6%) nor publication bias was observed in these peaks (Funnel plots in Supplementary Material 3). However, none of these brain regions survived correction for multiple testing. After including PD medication dose, expressed LEDD, as a covariate, grey matter reduction in right precuneus remained significant in PDP patients compared to PDnP patients (extending to left precuneus, and bilateral cuneus; voxel number = 1084, *Z* = −2.952, *p* = 0.001). There was substantial overlap between previously identified peaks, with the largest clusters in the left angular gyrus (extending to left inferior parietal gyrus; voxel number = 389, *Z*=-2.596, *p* = 0.004), right inferior temporal gyrus (extending to right middle temporal gyrus and right fusiform; voxel number = 353, *Z* =-.2921, *p* = 0.001), the left middle occipital gyrus (extending to inferior occipital gyrus; voxel number = 315, *Z*=-2.902, *p* = 0.001), and the right inferior parietal gyrus (extending to right angular gyrus; voxel number = 304, *Z*=-2.525, *p* = 0.005) (eFigure2, Supplementary Material 3). No publication bias was observed in any of these analyses (Funnel plots in Supplementary Material 3). However, none of these brain regions survived correction for multiple testing. When cognitive scores were entered as covariate, the right precuneus remained the largest cluster showing grey matter volume reduction (extending to the left precuneus, bilateral cuneus, voxel number = 1186, *Z* = −3.287, *p* < 0.001). In addition, this analysis showed other areas of lower grey matter volume in PDP compared to PDnP patients were observed. These were the left inferior occipital gyrus (extending to the middle occipital gyrus and left optic radiation, voxel number = 252, *Z* = −2.508, *p* = 0.006), the left postcentral gyrus (extending to the left precuneus, voxel number = 116, *Z* = −2.358, *p* = 0.009), and the right superior parietal gyrus (voxel number = 28, *Z* = −2.030, *p* = 0.021) (eFigure3, Supplementary Material 3). No publication bias was observed in any of these analyses (Funnel plots in Supplementary Material 3). The overlap between brain regions across the unadjusted and adjusted analyses is shown in Figure 2, and Table 2 reports peaks coordinates from the three analyses. We did not observe any significant peak in the opposite direction (i.e., ‘PDP > PDnP’) in either LEDD- or cognitive score-adjusted analyses. None of the areas across the three analyses survived familywise error correction.

**Figure 2.**
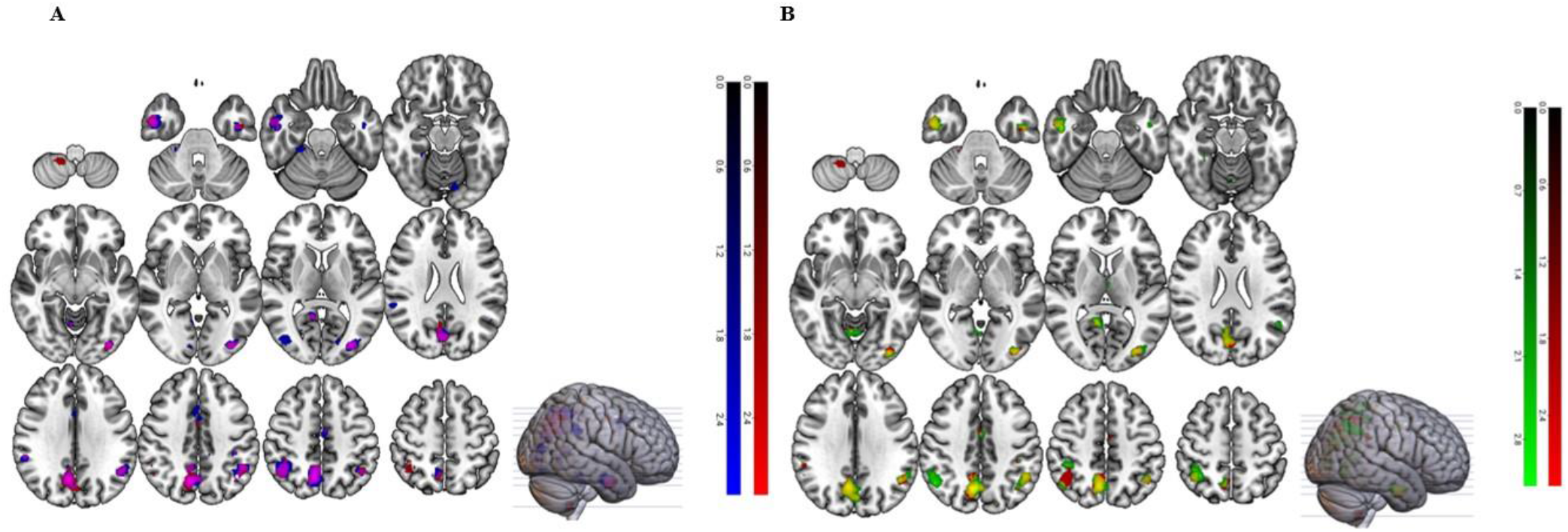
A) Overlapping peak areas (shown in purple) with grey matter loss in PD psychosis patients unadjusted (red) and adjusted for LEDD (expressed in mg/day) (blue) as covariate (uncorrected). This is indicated by the red and blue colour bars on which represent the T threshold of the voxels within the map. B) Overlapping peak areas (shown in yellow) with grey matter loss in PD psychosis patients unadjusted (red) and adjusted for cognitive scores (green) as covariate (uncorrected). This is indicated by the red and green colours bar on the right-hand side of the figure which represent the T threshold of the voxels within the map. The left side of the brain is shown on the right side of these brain images.

**Table 2.**
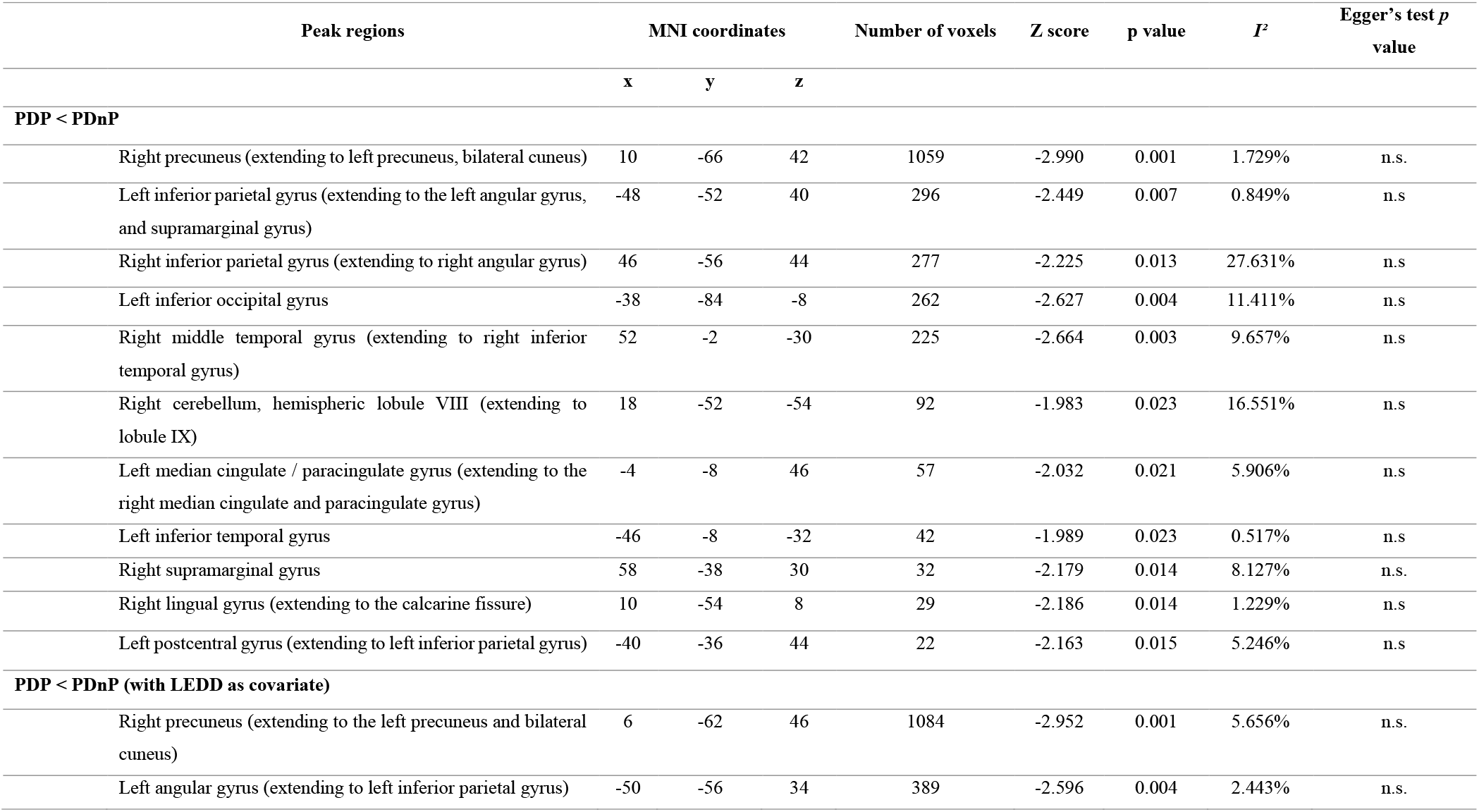
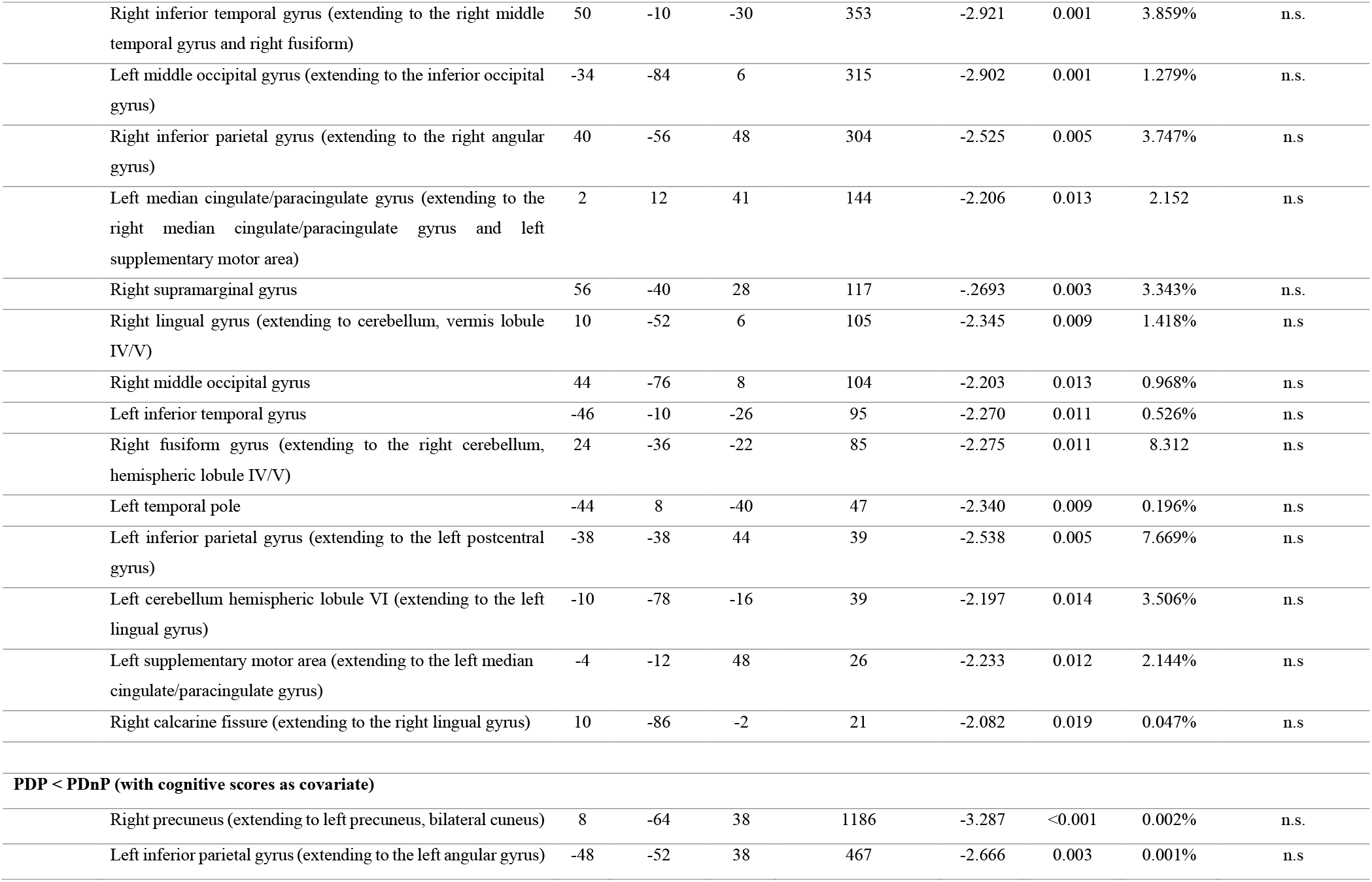
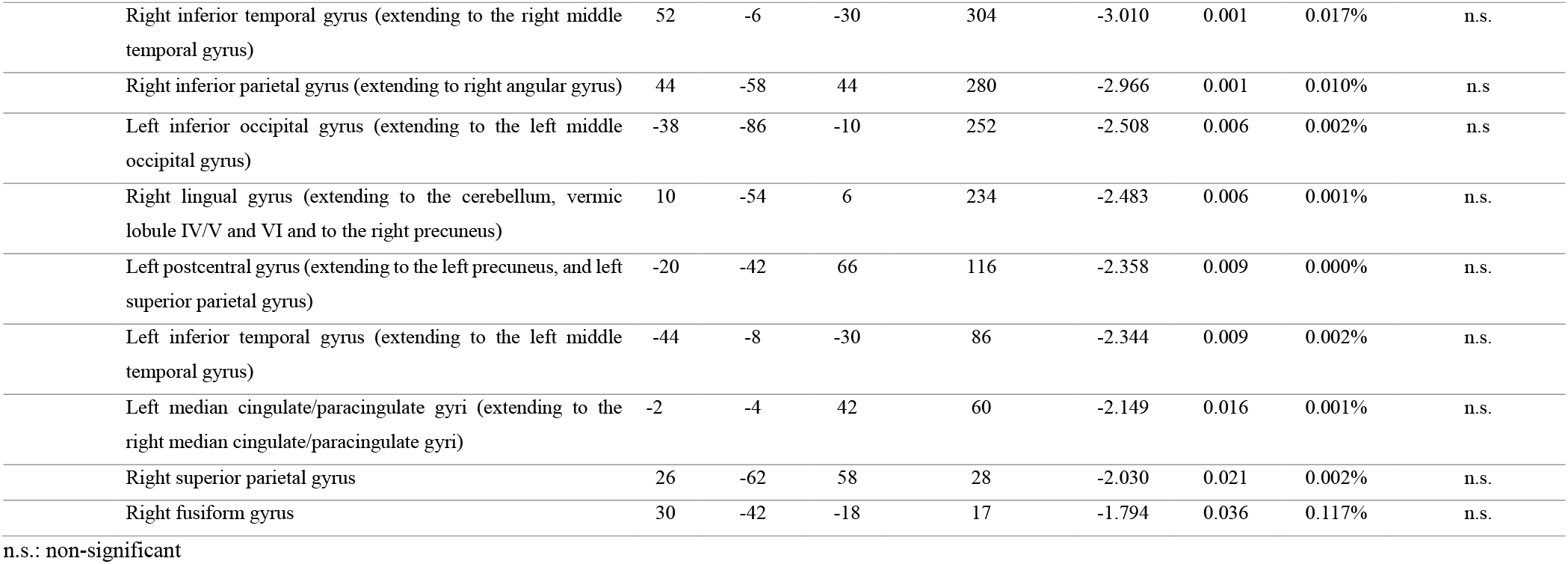
Peak coordinates showing greater grey matter loss in PDP patients compared to PDnP patients, alongside number of voxels, Z scores, and associated *p* value (uncorrected), *I^2^* (which indicates the magnitude of between-study heterogeneity as a proportion of total variability within each peak), and publication bias expressed as Egger’s test *p* value.

Neuroanatomical substrates in PD psychosis
| Peak regions | MNI coordinates | | | Number of voxels | Z score | p value | $I^2$ | Egger's test $p$ value |
| --- | --- | --- | --- | --- | --- | --- | --- | --- |
|  | x | y | z |  |  |  |  |  |
| PDP < PDnP |  |  |  |  |  |  |  |  |
| Right precuneus (extending to left precuneus, bilateral cuneus) | 10 | -66 | 42 | 1059 | -2.990 | 0.001 | 1.729% | n.s. |
| Left inferior parietal gyrus (extending to the left angular gyrus, and supramarginal gyrus) | -48 | -52 | 40 | 296 | -2.449 | 0.007 | 0.849% | n.s |
| Right inferior parietal gyrus (extending to right angular gyrus) | 46 | -56 | 44 | 277 | -2.225 | 0.013 | 27.631% | n.s |
| Left inferior occipital gyrus | -38 | -84 | -8 | 262 | -2.627 | 0.004 | 11.411% | n.s |
| Right middle temporal gyrus (extending to right inferior temporal gyrus) | 52 | -2 | -30 | 225 | -2.664 | 0.003 | 9.657% | n.s |
| Right cerebellum, hemispheric lobule VIII (extending to lobule IX) | 18 | -52 | -54 | 92 | -1.983 | 0.023 | 16.551% | n.s |
| Left median cingulate / paracingulate gyrus (extending to the right median cingulate and paracingulate gyrus) | -4 | -8 | 46 | 57 | -2.032 | 0.021 | 5.906% | n.s |
| Left inferior temporal gyrus | -46 | -8 | -32 | 42 | -1.989 | 0.023 | 0.517% | n.s |
| Right supramarginal gyrus | 58 | -38 | 30 | 32 | -2.179 | 0.014 | 8.127% | n.s. |
| Right lingual gyrus (extending to the calcarine fissure) | 10 | -54 | 8 | 29 | -2.186 | 0.014 | 1.229% | n.s |
| Left postcentral gyrus (extending to left inferior parietal gyrus) | -40 | -36 | 44 | 22 | -2.163 | 0.015 | 5.246% | n.s |
| PDP < PDnP (with LEDD as covariate) |  |  |  |  |  |  |  |  |
| Right precuneus (extending to the left precuneus and bilateral cuneus) | 6 | -62 | 46 | 1084 | -2.952 | 0.001 | 5.656% | n.s. |
| Left angular gyrus (extending to left inferior parietal gyrus) | -50 | -56 | 34 | 389 | -2.596 | 0.004 | 2.443% | n.s. |
| Right inferior temporal gyrus (extending to the right middle temporal gyrus and right fusiform) | 50 | -10 | -30 | 353 | -2.921 | 0.001 | 3.859% | n.s. |
| Left middle occipital gyrus (extending to the inferior occipital gyrus) | -34 | -84 | 6 | 315 | -2.902 | 0.001 | 1.279% | n.s. |
| Right inferior parietal gyrus (extending to the right angular gyrus) | 40 | -56 | 48 | 304 | -2.525 | 0.005 | 3.747% | n.s. |
| Left median cingulate/paracingulate gyrus (extending to the right median cingulate/paracingulate gyrus and left supplementary motor area) | 2 | 12 | 41 | 144 | -2.206 | 0.013 | 2.152 | n.s. |
| Right supramarginal gyrus | 56 | -40 | 28 | 117 | -.2693 | 0.003 | 3.343% | n.s. |
| Right lingual gyrus (extending to cerebellum, vermis lobule IV/V) | 10 | -52 | 6 | 105 | -2.345 | 0.009 | 1.418% | n.s. |
| Right middle occipital gyrus | 44 | -76 | 8 | 104 | -2.203 | 0.013 | 0.968% | n.s. |
| Left inferior temporal gyrus | -46 | -10 | -26 | 95 | -2.270 | 0.011 | 0.526% | n.s. |
| Right fusiform gyrus (extending to the right cerebellum, hemispheric lobule IV/V) | 24 | -36 | -22 | 85 | -2.275 | 0.011 | 8.312 | n.s. |
| Left temporal pole | -44 | 8 | -40 | 47 | -2.340 | 0.009 | 0.196% | n.s. |
| Left inferior parietal gyrus (extending to the left postcentral gyrus) | -38 | -38 | 44 | 39 | -2.538 | 0.005 | 7.669% | n.s. |
| Left cerebellum hemispheric lobule VI (extending to the left lingual gyrus) | -10 | -78 | -16 | 39 | -2.197 | 0.014 | 3.506% | n.s. |
| Left supplementary motor area (extending to the left median cingulate/paracingulate gyrus) | -4 | -12 | 48 | 26 | -2.233 | 0.012 | 2.144% | n.s. |
| Right calcarine fissure (extending to the right lingual gyrus) | 10 | -86 | -2 | 21 | -2.082 | 0.019 | 0.047% | n.s. |
| **PDP < PDnP (with cognitive scores as covariate)** |  |  |  |  |  |  |  |  |
| Right precuneus (extending to left precuneus, bilateral cuneus) | 8 | -64 | 38 | 1186 | -3.287 | <0.001 | 0.002% | n.s. |
| Left inferior parietal gyrus (extending to the left angular gyrus) | -48 | -52 | 38 | 467 | -2.666 | 0.003 | 0.001% | n.s. |
| Right inferior temporal gyrus (extending to the right middle temporal gyrus) | 52 | -6 | -30 | 304 | -3.010 | 0.001 | 0.017% | n.s. |
| Right inferior parietal gyrus (extending to right angular gyrus) | 44 | -58 | 44 | 280 | -2.966 | 0.001 | 0.010% | n.s. |
| Left inferior occipital gyrus (extending to the left middle occipital gyrus) | -38 | -86 | -10 | 252 | -2.508 | 0.006 | 0.002% | n.s. |
| Right lingual gyrus (extending to the cerebellum, vermic lobule IV/V and VI and to the right precuneus) | 10 | -54 | 6 | 234 | -2.483 | 0.006 | 0.001% | n.s. |
| Left postcentral gyrus (extending to the left precuneus, and left superior parietal gyrus) | -20 | -42 | 66 | 116 | -2.358 | 0.009 | 0.000% | n.s. |
| Left inferior temporal gyrus (extending to the left middle temporal gyrus) | -44 | -8 | -30 | 86 | -2.344 | 0.009 | 0.002% | n.s. |
| Left median cingulate/paracingulate gyri (extending to the right median cingulate/paracingulate gyri) | -2 | -4 | 42 | 60 | -2.149 | 0.016 | 0.001% | n.s. |
| Right superior parietal gyrus | 26 | -62 | 58 | 28 | -2.030 | 0.021 | 0.002% | n.s. |
| Right fusiform gyrus | 30 | -42 | -18 | 17 | -1.794 | 0.036 | 0.117% | n.s. |

n.s.: non-significant

### 3.3. Whole brain correlations with D1/D2 and 5-HT2a/5-HT1a gene expressions

Due to presence of multicollinearity in the multiple linear regression model, D2 receptor data were dropped from the analysis (i.e., due to a variation inflation factor, VIF = 6.5). Multiple linear regression analysis showed a significant association between Hedges’ *g* effect-size estimates of grey matter volume unadjusted for LEDD and 5-HT2a gene expression (regression coefficient =-0.107 (95% CI, −0.174, −0.039), t=-3.147, *p* = 0.002) and 5-HT1a gene expression (regression coefficient = 0.109 (95% CI, 0.024, 0.193), t = 2.565, *p* = 0.012) but not with D1 gene expression (D1, *p* = 0.554) across the 78 regions of the Desikan-Killiany atlas. Separate analysis with Hedges’ *g* effect-size estimates for grey matter volume adjusted for LEDD did not change the result (5-HT2a, regression coefficient =-0.120, 95% CI −0.190, −0.050, t=-3.408, *p*=0.001; 5-HT1a, regression coefficient =0.126, 95% CI 0.038, 0.213, t=2.848, *p*=0.006; D1, *p*=0.597). Presence of multicollinearity was also detected in this model and D2 was removed (VIF=6.67). Figure 3 reports the association between grey matter volume loss and receptor density from the LEDD-adjusted analysis. In both multiple linear regression models (adjusted and unadjusted for LEDD), the associations between regional cortical volume and serotonergic gene expressions were consistent in terms of direction of relationship. However, these associations were in opposite directions for the two serotonergic receptors, whereby the less the grey matter volume the more the 5-HT2a gene expression density and the less the 5-HT1a gene expression density across the 78 regions of the Desikan-Killiany atlas. That is, the greater the grey matter reduction the greater the 5-HT2a and the lower 5-HT1a gene expression density. Figures showing the relationship between cortical and subcortical spatial gene expressions and Hedges’ *g* effect sizes unadjusted for LEDD are in Supplementary Material 4 (eFigure4-5).

**Figure 3.**
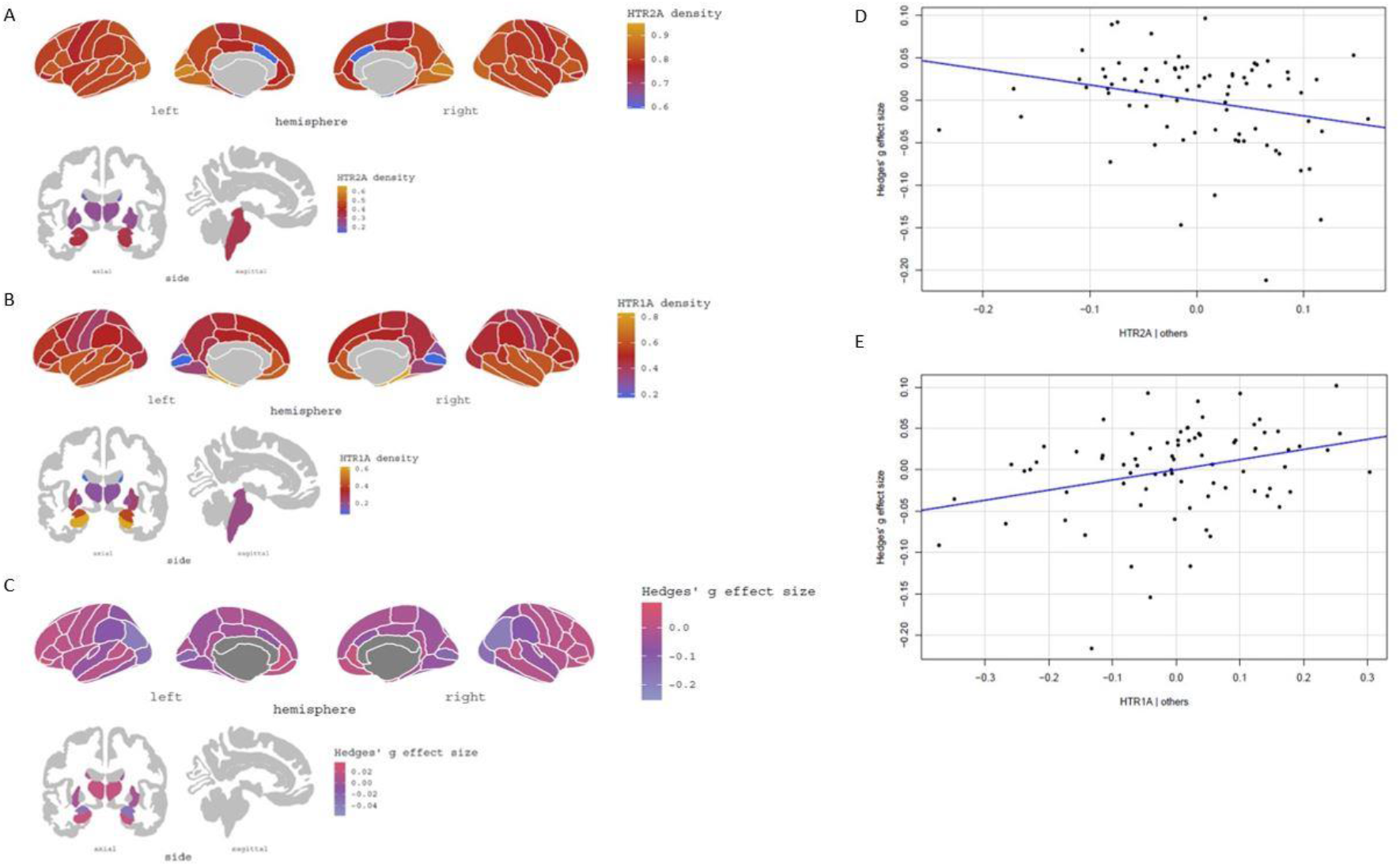
Gene expression density of 5-HT2a (A), 5-HT1a (B) receptors in cortical and subcortical regions, and Hedges’ g effect size in cortical and subcortical regions (C) derived from the covariate meta-analysis (adjusted for LEDD) results showing decreased grey matter in PDP patients compared to PDnP patients (PDP < PDnP), parcellated across the 78 brain regions of the Desikan-Killiany atlas (Desikan et al., 2006). On the right-hand side, scatterplots showing the relationship between 5-HT2a (D) and 5-HT1a (E) with Hedges’ g effect-size estimates of grey matter volume adjusted for LEDD in PDP patients compared to PDnP patients (the regression lines are adjusted for all the predictors).

## 4. Discussion

Findings from individual studies reported extensive areas of lower grey matter volume in PDP without providing clear understanding on whether PDP may be due to structural issues in key regions or diffused anomalies. Here, we have addressed this ambiguity by incorporating results from structural MRI studies using a quantitative approach to identify neuroanatomical correlates implicated in PDP. The meta-analytic results found widespread grey matter volume reduction across parietal-temporal-occipital regions in PDP patients, specifically, the right precuneus (extending to the left precuneus), bilateral inferior parietal gyrus, and left inferior occipital gyrus, corroborating previous evidence (Goldman et al., 2014; Lenka et al., 2015; Yao et al., 2014). Volume reduction in right precuneus, bilateral inferior parietal gyrus, left median cingulate/paracingulate cortex, and right inferior and middle temporal gyrus remained significant and was independent of PD medications and cognitive scores. When PD medications and cognitive scores standardised across studies were included as a covariate, there were additional areas of lower cortical volume within occipital (e.g., left middle occipital gyrus), parietal (e.g., right lingual and left angular gyrus), and temporal regions (e.g., right fusiform gyrus). These results also extend on previous evidence (Bejr-Kasem et al., 2019; Jia et al., 2019; Lenka et al., 2015; Yao et al., 2014). Across the analyses, we identified large clusters of grey matter loss in regions associated with higher order visual processing (i.e., dorsal and ventral visual pathways) and the Default Mode Network (DMN), and specifically the precuneus, one of the DMN nodes, which is embedded in the parietal lobe and is involved in information processing (including visual information) (Buckner et al., 2008; Cavanna & Trimble, 2006; Freton et al., 2014; Hafkemeijer et al., 2012; Lenka et al., 2015). Grey matter volume loss in these areas may lead to dysfunctional information integration giving rise to visual hallucinations in PD patients, and also potentially due to an overreliance on endogenous attentional mechanisms (Shine et al., 2011; Shine, O’Callaghan, et al., 2014) as suggested by the involvement of DMN nodes (Rektorova et al., 2014).

Furthermore, this review expands on previous evidence by investigating the association between regional grey matter volume (as measured by Hedges’ *g* effect-size estimates from the meta-analysis) and key candidate receptors involved in PDP as indexed by mRNA microarray gene expression extracted from the Allen Human Brain Atlas (Hawrylycz et al., 2012). Expression of 5-HT2a and 5-HT1a receptors were associated with estimates of grey matter volume, albeit being in opposite direction. Whether the inverse relationship observed here between pooled grey matter volume in PDP patients and the pooled expression of 5-HT2a receptors in an independent healthy cohort indicates that in PDP patients the greater the grey matter volume loss the greater is the regional 5-HT2a receptor availability, remains to be tested by direct investigation using both imaging modalities in the same group of participants. Nevertheless, if an association between grey matter loss in PDP and increased 5-HT2a receptor availability is demonstrated in PDP, it would be consistent with pharmacological evidence of the efficacy of the atypical antipsychotic Pimavanserin, a highly selective 5-HT2a receptor antagonist, in the treatment of psychosis in PD (Majlath et al., 2017; Meltzer et al., 2010; Mohanty et al., 2019; Stahl, 2016a; Yunusa et al., 2020). On the other hand, the direct relationship between grey matter volume and the pooled expression of 5-HT1a receptors, may indicate that in PDP patients the lower the grey matter volume the lower the regional availability of 5-HT1a receptors. Whether this holds true when investigated using both imaging modalities in the same cohort of patients remains to be seen. Given the association between 5-HT1a abnormalities and depression (Wang et al., 2016), this may reflect the common co-occurrence of depressive symptoms in people with PDP. While the degeneration of dopaminergic neurons in the substantia nigra is the hallmark of PD pathology, it is also followed by degeneration of serotonergic neurons in subcortical areas such as the raphe nuclei (Stahl, 2016b). Further, upregulation of 5-HT2a receptors has been observed in visual and temporal regions in PDP patients which may be a result of Lewy bodies accumulation in subcortical regions which also was associated with presence of hallucinatory behaviour (Birkmayer et al., 1974; Huot, 2018; Huot et al., 2010). Our results are in line with those from Vignando et al.’s (Vignando et al., 2021) analysis showing association between structural measures and 5-HT2a and 5-HT1a binding but not in terms of relationship with regional D1 density.

Presumably, the development of psychosis in PD patients could then be due to dysfunction in networks involved in attention control and visual information processing (Lenka et al., 2015; Muller et al., 2014; Shine, Halliday, et al., 2014; Shine et al., 2011; Shine, O’Callaghan, et al., 2014) that become pathological due to grey matter reduction in their nodes, as shown in the meta-analysis results, which consequently leads to abnormal functionality at a network level. Our results are to some extent different from those of the mega-analysis (Vignando et al., 2021) and may be due to the diverse methodologies used (mega-analysis vs. meta-analysis), the qualitative synthesis of multi-modal neuroimaging studies, the parameters used to assess atrophy (cortical thickness and surface area vs. voxel-based grey matter), and the availability of raw data (subject-level data vs. peak coordinates and one T map). They reported reduced surface area and extensive reduction in cortical thickness in occipital, temporal, parietal, frontal and limbic regions in PDP patients with visual hallucinations, identifying asymmetry in the left ventral visual pathway and extensive cortical thinning in bilateral cuneus, left dorso-medial superior frontal gyrus. Conversely, we mainly reported parietal-temporo-occipital regions as areas of significant grey matter loss and two frontal regions were found involved in PDP (the latter observed in the main analysis and in the analysis adjusted for cognitive state), our results however did not survive familywise error correction. Interestingly, we identified the right precuneus (extending to the left precuneus) as the largest cluster affected by grey matter loss in PDP patients across the three analyses (i.e., unadjusted, adjusted for PD medication and for cognition). The cuneus and precuneus were also reported as two of the areas with the greatest effect size in the principal component analysis by Vignando et al. (2021) examining the nodes that contributed the most to cortical thickness reduction in PDP patients, as well as in their network analysis. We conducted a whole-brain association between grey matter loss in PDP patients and receptor density examining dopaminergic and serotonergic receptors. We did not observe any relationships between grey matter volume and regional D1 density. This is in contrast with Vignando et al. (2021) who found associations with dopaminergic receptor density with cortical thickness in regions where PDP and PDnP had significant differences, and with surface area for significant regions of difference as well as across the cortex. However, we were only able to extract data on D1/D2 receptors unlike Vignando et al. (2021) who investigated the relationship with D2/D3 joint receptor density distribution. This discrepancy may also be due to methodological differences: they used mean thickness differences as the outcome variable, whilst we employed Hedges’ *g* effect-size estimates extracted from SDM-PSI used as a measure of grey matter loss in PDP patients, lastly they examined in-vivo PET data from an independent healthy cohort whilst we relied on gene expression data extracted from six healthy donors of the Allen Human Brain Atlas (Arnatkevičiūtė et al., 2019; Desikan et al., 2006).

### 4.1. Strength and limitations

The main limitations were the lack of significant results in the family-wise error corrected analyses potentially due to the small number studies and the limited access to raw imaging data as we mainly relied on peak coordinates, a less powerful approach (Salimi-Khorshidi et al., 2009), and that we have fewer studies than recommended guidelines (Müller et al., 2018). We were also unable to relate meta-analytic estimates with psychotic symptoms. These studies focused on visual hallucinations in PD. This may also be due to the different rating scales applied to assess the presence of these symptoms, some measures may not differentiate or may not be sensitive to these symptoms. Whilst some studies referred to the NINDS-NIMH criteria (Ravina et al., 2007), most used different assessment tools or interviews. Although visual hallucinations and illusions are the most prevalent psychotic manifestations in PD, there is evidence of patients experiencing auditory and multimodal (e.g., olfactory, tactile (Chou et al., 2005; Solla et al., 2021)) hallucinations and delusions (Goetz et al., 2006; Marsh et al., 2004; Papapetropoulos et al., 2008) as the disease progresses, leading to increased distress and worsening of quality of life for the patients. Future exploration should compare the neuroanatomical correlates of different psychotic symptoms in PD, as this can shed light on whether there are common pathways shared with visual hallucinations, as those observed in this meta-analysis, or symptom-specific dysfunction. A further limitation is that we used mRNA expression, a result of transcriptional and translation activity, as a measure of dopaminergic (i.e., D1, D2) and serotonergic (i.e., 5-HT2a, 5-HT1a) gene expression extracted from 6 neurotypical donors (Hawrylycz et al., 2012) which may not be presentative of the PD population and may introduce variability due to inter-individual differences. Nevertheless, several studies employing this methodology have shown the consistency of the receptor spatial architecture in the cortex across individuals (Beliveau et al., 2017; Rizzo et al., 2016; Rizzo et al., 2014; Selvaggi et al., 2019; Veronese et al., 2016). Similar approaches have been applied by other studies assessing the relationship between estimates of brain functions (from fMRI studies) and regional gene expression data, also under pharmacological intervention (Anderson et al., 2020; Gryglewski et al., 2018; Richiardi et al., 2015; Selvaggi et al., 2019; Vértes et al.). Using mRNA expression data can pave the way in the use of different techniques, also more available to the wider community, to measure regional receptor expression in the whole brain when there is a paucity of in-vivo data.

### 4.2. Conclusions

In conclusion, there was grey matter volume loss in the parietal-temporal-occipital region which persisted even after adjusting for the effects of PD medications and cognitive scores. Although it may be premature to infer definite conclusions, the above evidence suggests that anomalies in brain regions involved in processing visual stimuli, over reliance on internal processing, and potential deficits in directing attention and integrating endogenous and externally derived information may underlie psychosis in PD. These findings relate to PD patients with mainly visual hallucinations due to the predominance of this symptom in the cohorts included in these studies. Furthermore, we observed an association between regional gene expression of serotonergic receptors and regional grey matter volume.

## Acknowledgment

None

## Authors’ contributions

*Conceptualisation:* Sara Pisani, Latha Velayudhan, Sagnik Bhattacharyya

*Methodology:* Sara Pisani, Brandon Gunasekera, Latha Velayudhan, Sagnik Bhattacharyya

*Investigation:* Sara Pisani, Latha Velayudhan, Sagnik Bhattacharyya

*Data curation:* Sara Pisani, Yining Lu

*Formal analysis:* Sara Pisani, Brandon Gunasekera, Yining Lu, Sagnik Bhattacharyya

*Visualisation:* Sarra Pisani, Sagnik Bhattacharyya

*Funding acquisition*: Sagnik Bhattacharyya, Latha Velayudhan, Dominic ffytche, Dag

Aarsland, Kallol Ray Chaudhuri, Clive Ballard

*Writing (original draft):* Sara Pisani, Sagnik Bhattacharyya, Latha Velayudhan

*Writing (review & editing):* All authors

*Supervision:* Latha Velayudhan, Dominic ffytche, Sagnik Bhattacharyya

