## Supplementary Material for "Neuroanatomical substrates in Parkinson’s Disease psychosis and their association with serotonergic receptor gene expression: A coordinate-based meta-regression analysis"

### Supplementary Material 1

Search strategy conducted on 7th June 2021 on Embase (Ovid)

1. Brain.mp. [mp=title, abstract, heading word, drug trade name, original title, device manufacturer, drug manufacturer, device trade name, keyword, floating subheading word, candidate term word]

2. Brain region*.mp. [mp=title, abstract, heading word, drug trade name, original title, device manufacturer, drug manufacturer, device trade name, keyword, floating subheading word, candidate term word]

3. Brain activit*.mp. [mp=title, abstract, heading word, drug trade name, original title, device manufacturer, drug manufacturer, device trade name, keyword, floating subheading word, candidate term word]

4. Functional connect*.mp. [mp=title, abstract, heading word, drug trade name, original title, device manufacturer, drug manufacturer, device trade name, keyword, floating subheading word, candidate term word]

5. Functional imaging.mp. [mp=title, abstract, heading word, drug trade name, original title, device manufacturer, drug manufacturer, device trade name, keyword, floating subheading word, candidate term word]

6. Neurophysiological mechanism*.mp. [mp=title, abstract, heading word, drug trade name, original title, device manufacturer, drug manufacturer, device trade name, keyword, floating subheading word, candidate term word]

7. Neuroanatomical correlate*.mp. [mp=title, abstract, heading word, drug trade name, original title, device manufacturer, drug manufacturer, device trade name, keyword, floating subheading word, candidate term word]

8. Neural substrate*.mp. [mp=title, abstract, heading word, drug trade name, original title, device manufacturer, drug manufacturer, device trade name, keyword, floating subheading word, candidate term word]

9. Neural correlate*.mp. [mp=title, abstract, heading word, drug trade name, original title, device manufacturer, drug manufacturer, device trade name, keyword, floating subheading word, candidate term word]

10. Cerebral mechanism*.mp. [mp=title, abstract, heading word, drug trade name, original title, device manufacturer, drug manufacturer, device trade name, keyword, floating subheading word, candidate term word]

11. Cerebral atrophy.mp. [mp=title, abstract, heading word, drug trade name, original title, device manufacturer, drug manufacturer, device trade name, keyword, floating subheading word, candidate term word]

12. Whole brain analys*.mp. [mp=title, abstract, heading word, drug trade name, original title, device manufacturer, drug manufacturer, device trade name, keyword, floating subheading word, candidate term word]

13. "MRI".mp. [mp=title, abstract, heading word, drug trade name, original title, device manufacturer, drug manufacturer, device trade name, keyword, floating subheading word, candidate term word]

14. Magnetic resonance imaging.mp. [mp=title, abstract, heading word, drug trade name, original title, device manufacturer, drug manufacturer, device trade name, keyword, floating subheading word, candidate term word]

15. Structural MRI.mp. [mp=title, abstract, heading word, drug trade name, original title, device manufacturer, drug manufacturer, device trade name, keyword, floating subheading word, candidate term word]

16. Structural magnetic resonance imaging.mp. [mp=title, abstract, heading word, drug trade name, original title, device manufacturer, drug manufacturer, device trade name, keyword, floating subheading word, candidate term word]

17. "fMRI".mp. [mp=title, abstract, heading word, drug trade name, original title, device manufacturer, drug manufacturer, device trade name, keyword, floating subheading word, candidate term word]

18. Functional MRI.mp. [mp=title, abstract, heading word, drug trade name, original title, device manufacturer, drug manufacturer, device trade name, keyword, floating subheading word, candidate term word]

19. Functional magnetic resonance imaging.mp. [mp=title, abstract, heading word, drug trade name, original title, device manufacturer, drug manufacturer, device trade name, keyword, floating subheading word, candidate term word]

20. Resting state fMRI.mp. [mp=title, abstract, heading word, drug trade name, original title, device manufacturer, drug manufacturer, device trade name, keyword, floating subheading word, candidate term word]

21. "rsfMRI".mp. [mp=title, abstract, heading word, drug trade name, original title, device manufacturer, drug manufacturer, device trade name, keyword, floating subheading word, candidate term word]

22. tractography.mp. [mp=title, abstract, heading word, drug trade name, original title, device manufacturer, drug manufacturer, device trade name, keyword, floating subheading word, candidate term word]

23. Diffusion tensor imaging.mp. [mp=title, abstract, heading word, drug trade name, original title, device manufacturer, drug manufacturer, device trade name, keyword, floating subheading word, candidate term word]

24. "DTI".mp. [mp=title, abstract, heading word, drug trade name, original title, device manufacturer, drug manufacturer, device trade name, keyword, floating subheading word, candidate term word]

25. Positron emission tomography.mp. [mp=title, abstract, heading word, drug trade name, original title, device manufacturer, drug manufacturer, device trade name, keyword, floating subheading word, candidate term word]

26. "PET".mp. [mp=title, abstract, heading word, drug trade name, original title, device manufacturer, drug manufacturer, device trade name, keyword, floating subheading word, candidate term word]

27. "SPECT".mp. [mp=title, abstract, heading word, drug trade name, original title, device manufacturer, drug manufacturer, device trade name, keyword, floating subheading word, candidate term word]

28. Single photon emission computed tomography.mp. [mp=title, abstract, heading word, drug trade name, original title, device manufacturer, drug manufacturer, device trade name, keyword, floating subheading word, candidate term word]

29. arterial spin labelling.mp. [mp=title, abstract, heading word, drug trade name, original title, device manufacturer, drug manufacturer, device trade name, keyword, floating subheading word, candidate term word]

30. Voxel-base*.mp. [mp=title, abstract, heading word, drug trade name, original title, device manufacturer, drug manufacturer, device trade name, keyword, floating subheading word, candidate term word]

31. Voxel-base morphometry.mp. [mp=title, abstract, heading word, drug trade name, original title, device manufacturer, drug manufacturer, device trade name, keyword, floating subheading word, candidate term word]

32. "VBM".mp. [mp=title, abstract, heading word, drug trade name, original title, device manufacturer, drug manufacturer, device trade name, keyword, floating subheading word, candidate term word]

33. Magnetic resonance spectroscopy.mp. [mp=title, abstract, heading word, drug trade name, original title, device manufacturer, drug manufacturer, device trade name, keyword, floating subheading word, candidate term word]

34. "MRS".mp. [mp=title, abstract, heading word, drug trade name, original title, device manufacturer, drug manufacturer, device trade name, keyword, floating subheading word, candidate term word]

35. Neuroimaging.mp. [mp=title, abstract, heading word, drug trade name, original title, device manufacturer, drug manufacturer, device trade name, keyword, floating subheading word, candidate term word]

36. 1 or 2 or 3 or 4 or 5 or 6 or 7 or 8 or 9 or 10 or 11 or 12 or 13 or 14 or 15 or 16 or 17 or 18 or 19 or 20 or 21 or 22 or 23 or 24 or 25 or 26 or 27 or 28 or 29 or 30 or 31 or 32 or 33 or 34 or 35

37. Parkinson's disease psychosis.mp. [mp=title, abstract, heading word, drug trade name, original title, device manufacturer, drug manufacturer, device trade name, keyword, floating subheading word, candidate term word]

38. Parkinson disease psychosis.mp. [mp=title, abstract, heading word, drug trade name, original title, device manufacturer, drug manufacturer, device trade name, keyword, floating subheading word, candidate term word]

39. 37 or 38

40. exp Parkinson disease/

41. Parkinson*.mp. [mp=title, abstract, heading word, drug trade name, original title, device manufacturer, drug manufacturer, device trade name, keyword, floating subheading word, candidate term word]

42. Parkinsonian.mp. [mp=title, abstract, heading word, drug trade name, original title, device manufacturer, drug manufacturer, device trade name, keyword, floating subheading word, candidate term word]

43. Parkinsonism.mp. [mp=title, abstract, heading word, drug trade name, original title, device manufacturer, drug manufacturer, device trade name, keyword, floating subheading word, candidate term word]

44. Atypical parkinsonism.mp. [mp=title, abstract, heading word, drug trade name, original title, device manufacturer, drug manufacturer, device trade name, keyword, floating subheading word, candidate term word]

45. 40 or 41 or 42 or 43 or 44

46. exp psychosis/

47. Psychotic.mp. [mp=title, abstract, heading word, drug trade name, original title, device manufacturer, drug manufacturer, device trade name, keyword, floating subheading word, candidate term word]

48. Psychotic disorder*.mp. [mp=title, abstract, heading word, drug trade name, original title, device manufacturer, drug manufacturer, device trade name, keyword, floating subheading word, candidate term word]

49. Paranoia.mp. [mp=title, abstract, heading word, drug trade name, original title, device manufacturer, drug manufacturer, device trade name, keyword, floating subheading word, candidate term word]

50. Paranoid.mp. [mp=title, abstract, heading word, drug trade name, original title, device manufacturer, drug manufacturer, device trade name, keyword, floating subheading word, candidate term word]

51. Delusi*.mp. [mp=title, abstract, heading word, drug trade name, original title, device manufacturer, drug manufacturer, device trade name, keyword, floating subheading word, candidate term word]

52. Halluci*.mp. [mp=title, abstract, heading word, drug trade name, original title, device manufacturer, drug manufacturer, device trade name, keyword, floating subheading word, candidate term word]

53. Visual halluci*.mp. [mp=title, abstract, heading word, drug trade name, original title, device manufacturer, drug manufacturer, device trade name, keyword, floating subheading word, candidate term word]

54. Auditory halluci*.mp. [mp=title, abstract, heading word, drug trade name, original title, device manufacturer, drug manufacturer, device trade name, keyword, floating subheading word, candidate term word]

55. multimodal halluci*.mp. [mp=title, abstract, heading word, drug trade name, original title, device manufacturer, drug manufacturer, device trade name, keyword, floating subheading word, candidate term word]

56. Visual illusion*.mp. [mp=title, abstract, heading word, drug trade name, original title, device manufacturer, drug manufacturer, device trade name, keyword, floating subheading word, candidate term word]

57. Illusion*.mp. [mp=title, abstract, heading word, drug trade name, original title, device manufacturer, drug manufacturer, device trade name, keyword, floating subheading word, candidate term word]

58. Imagery.mp. [mp=title, abstract, heading word, drug trade name, original title, device manufacturer, drug manufacturer, device trade name, keyword, floating subheading word, candidate term word]

59. Schizophrenia spectrum disorder*.mp. [mp=title, abstract, heading word, drug trade name, original title, device manufacturer, drug manufacturer, device trade name, keyword, floating subheading word, candidate term word]

60. Psychosis spectrum disorder*.mp. [mp=title, abstract, heading word, drug trade name, original title, device manufacturer, drug manufacturer, device trade name, keyword, floating subheading word, candidate term word]

61. 46 or 47 or 48 or 49 or 50 or 51 or 52 or 53 or 54 or 55 or 56 or 57 or 58 or 59 or 60

62. 45 and 61

63. 39 or 62

64. 36 and 63

Results, N = 6368

Search strategy conducted on 7th June 2021 on PubMed

| Search number | Query | Sort By | Filters | Results |
|---|---|---|---|---|
| 7 | (#1) AND (#6) | | | 1,289 |
| 6 | (#2) OR (#5) | | | 3,452 |
| 5 | (#3) AND (#4) | | | 3,128 |
| 4 | (((((((((((((Psychosis[MeSH Terms]) OR (Psychotic)) OR (Psychotic disorder*)) OR (Paranoia)) OR (Paranoid)) OR (Delusi*)) OR (Halluci*)) OR (Visual halluci*)) OR (Auditory halluci*)) OR (Multimodal halluci*)) OR (Visual illusion*)) OR (Illusion*)) OR (Imagery)) OR (Schizophrenia spectrum disorder*)) OR (Psychosis spectrum disorder*) | | | 109,488 |
| 3 | ((((Parkinson disease[MeSH Terms]) OR (Parkinson*)) OR (Parkinsonian)) OR (Parkinsonism)) OR (Atypical Parkinsonism) | | | 91,133 |
| 2 | (Parkinson's disease psychosis) OR (Parkinson disease psychosis) | | | 1,383 |
| 1 | ((((((((((((((((((((((((((((((((Brain) OR (Brain region*)) OR (Brain activit*)) OR (Functional connect*)) OR (Functional imaging)) OR (Neurophysiological mechanism*)) OR (Neuroanatomical correlate*)) OR (Neural substrate*)) OR (Neural correlate*)) OR (Cerebral mechanism*)) OR (Cerebral atrophy)) OR (Whole brain analys*)) OR (MRI)) OR (Magnetic resonance imaging)) OR (Structural MRI)) OR (Structural magnetic resonance imaging)) OR (fMRI)) OR (Functional MRI)) OR (Functional magnetic resonance imaging)) OR (Resting state fMRI)) OR (rsfMRI)) OR (tractography)) OR (Diffusion tensor imaging)) OR (DTI)) OR (Positron emission tomorgraphy)) OR (PET)) OR (SPECT)) OR (single photon emission computed tomography)) OR (arterial spin labelling)) OR (Voxel-base*)) OR (Voxel-base morphometry)) OR (VBM)) OR (magnetic resonance spectroscopy)) OR (MRS)) OR (Neuroimaging) | | | 1,819,516 |

Search strategy conducted on 7th June 2021 on Web of Science

| Set | Results | Save History / Create AlertOpen Saved History |
|---|---|---|
| # 9 | 5,397 | #8 OR #4<br>*Indexes=SCI-EXPANDED, SSCI, A&HCI, CPCI-S, CPCI-SSH, ESCI Timespan=All years* |
| # 8 | 5,397 | #6 AND #5 |

| | | |
|---|---|---|
| | | *Indexes=SCI-EXPANDED, SSCI, A&HCI, CPCI-S, CPCI-SSH, ESCI Timespan=All years* |
| # 7 | 42,130 | #3 AND #2 AND #1<br>*Indexes=SCI-EXPANDED, SSCI, A&HCI, CPCI-S, CPCI-SSH, ESCI Timespan=All years* |
| # 6 | 260,356 | **ALL FIELDS:** (Psychosis) *OR* **ALL FIELDS:** (Psychotic) *OR* **ALL FIELDS:** ( Psychotic disorder*) *OR* **ALL FIELDS:** (Paranoia) *OR* **ALL FIELDS:** (Paranoid) *OR* **AL L FIELDS:** (Delusi*) *OR* **ALL FIELDS:** (Halluci*) *OR* **ALL FIELDS:** (Visual halluci*) *OR* **ALL FIELDS:** (Auditory halluci*) *OR* **ALL FIELDS:** (Multimodal halluci*) *OR* **ALL FIELDS:** (Visual illusion*) *OR* **ALL FIELDS:** (Illusion*) *OR* **ALL FIELDS:** (Imagery) *OR* **ALL FIELDS:** (Schizophrenia spectrum disorder*) *OR* **ALL FIELDS:** (Psychosis spectrum disorder)<br>*Indexes=SCI-EXPANDED, SSCI, A&HCI, CPCI-S, CPCI-SSH, ESCI Timespan=All years* |
| # 5 | 215,274 | **ALL FIELDS:** (Parkinson disease) *OR* **ALL FIELDS:** (Parkinson*) *OR* **ALL FIELDS:** (Parkinsonian) *OR* **ALL FIELDS:** (Parkinsonism) *OR* **ALL FIELDS:** (Atypical parkinsonism)<br>*Indexes=SCI-EXPANDED, SSCI, A&HCI, CPCI-S, CPCI-SSH, ESCI Timespan=All years* |
| # 4 | 1,727 | **ALL FIELDS:** (Parkinson's disease psychosis) *OR* **ALL FIELDS:** (Parkinson disease psychosis)<br>*Indexes=SCI-EXPANDED, SSCI, A&HCI, CPCI-S, CPCI-SSH, ESCI Timespan=All years* |
| # 3 | 327,942 | **ALL FIELDS:** (Voxel-base*) *OR* **ALL FIELDS:** (Voxel-base morphometry) *OR* **ALL FIELDS:** (VBM) *OR* **ALL FIELDS:** (Magnetic resonance spectroscopy) *OR* **ALL FIELDS:** (MRS) *OR* **ALL FIELDS:** (Neuroimaging)<br>*Indexes=SCI-EXPANDED, SSCI, A&HCI, CPCI-S, CPCI-SSH, ESCI Timespan=All years* |
| # 2 | 552,341 | **ALL FIELDS:** (Structual magnetic resonance imaging) *OR* **ALL FIELDS:** (fMRI) *OR* **ALL FIELDS:** (Functional MRI) *OR* **ALL FIELDS:** (Functional magnetic resonance imaging) *OR* **ALL FIELDS:** (Resting state fMRI) *OR* **ALL FIELDS:** (rsfMRI) *OR* **ALL FIELDS:** (Tractography) *OR* **ALL FIELDS:** (DTI) *OR* **ALL FIELDS:** (Diffusion tensor imaging) *OR* **ALL FIELDS:** (Positron emission tomography) *OR* **ALL FIELDS:** (PET) *OR* **ALL FIELDS:** (SPECT) *OR* **ALL F IELDS:** (Single photon emission computed tomography) *OR* **ALL FIELDS:** (Arterial spin labelling) |

| | | |
|---|---|---|
| | | *Indexes=SCI-EXPANDED, SSCI, A&HCI, CPCI-S, CPCI-SSH, ESCI Timespan=All years* |
| # 1 | 2,316,195 | **ALL FIELDS:** (Brain*) *OR* **ALL FIELDS:** (Brain region*) *OR* **ALL FIELDS:** (Functional connect*) *OR* **ALL FIELDS:** (Functional imaging) *OR* **ALL FIELDS:** (Neurophysiological mechanism*) *OR* **ALL FIELDS:** (Neuroanatomical correlate*) *OR* **ALL FIELDS:** (Neural substrate*) *OR* **ALL FIELDS:** (Neural correlate*) *OR* **ALL FIELDS:** (Cerebral mechanism*) *OR* **ALL FIELDS:** (Cerebral atrophy) *OR* **ALL FIELDS:** (Whole brain analys*) *OR* **ALL FIELDS:** (MRI) *OR* **ALL FIELDS:** (Magnetic resonance imaging) *OR* **ALL FIELDS:** (Structural MRI)<br><br>*Indexes=SCI-EXPANDED, SSCI, A&HCI, CPCI-S, CPCI-SSH, ESCI Timespan=All years* |

### Supplementary Material 2

eTable1 shows the quality rating of the 10 studies included in the review. Study quality was assessed with the Newcastle-Ottawa Scale in the three methodological domains, i.e., Selection, Comparability, and Exposure. Studies were assigned a maximum of one star per item with the exception of Comparability (i.e., maximum of two stars).

| Study | Selection | | | | Comparability | Exposure | | |
|---|---|---|---|---|---|---|---|---|
| | Is the case definition adequate? | Representativeness of the cases | Selection of controls | Definition of Controls | Comparability of cases and controls on the basis of the design and analysis | Ascertainment of exposure | Same method of ascertainment for cases and controls | Non-response rate |
| Bejr-Kasem et al. (Bejr-Kasem et al., 2019) | * | | * | | ** | * | * | * |
| Lee et al. (Lee et al., 2017) | * | | * | | ** | * | * | * |
| Pagonabarraga et al. (Pagonabarraga et al., 2014) | * | | * | | ** | * | * | * |
| Lawn & ffytche (Lawn, 2021) | * | | * | * | ** | * | * | * |
| Ramirez-Ruiz et al. (Ramirez-Ruiz et al., 2007) | * | | * | | * | * | * | * |
| Bejr-Kasem et al. (Bejr-kasem et al., 2021) | * | | * | | ** | * | * | * |

| | | | | | | | |
|---|---|---|---|---|---|---|---|
| **Watanabe et al. (Watanabe et al., 2013)** | * | * | * | ** | * | * | * |
| **Firbank et al. (Firbank et al., 2018)** | * | | * | ** | * | * | * |
| **Shin et al. (Shin et al., 2012)** | * | | * | ** | * | * | * |
| **Goldman et al. (Goldman et al., 2014)** | * | | * | ** | * | * | * |

### Supplementary Material 3

**eFigure1.** Peak areas with grey matter loss in PDP patients (uncorrected) compared to PDnP patients. This is indicated by the red colour bar on which represent the T threshold of the voxels within the map.

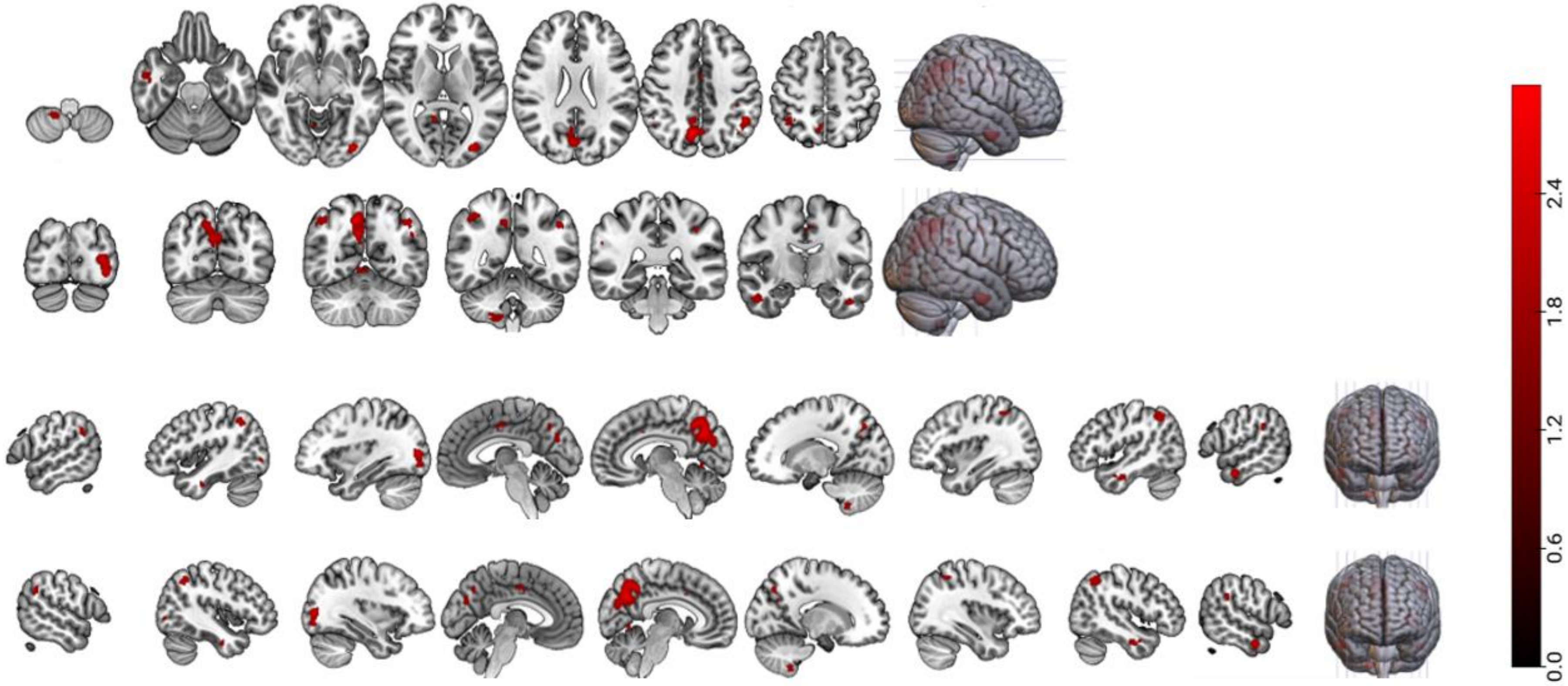

Funnel plots for each peak region (with *p* value) with significant grey matter volume loss in PDP patients compared to PDnP patients.

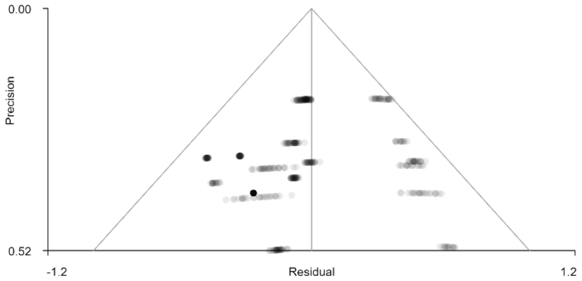

Right precuneus, p=0.600

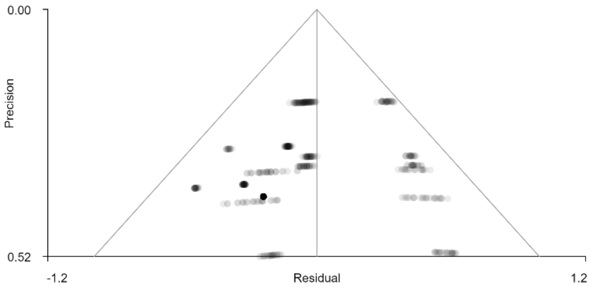

Left inferior parietal gyrus, p=0.768

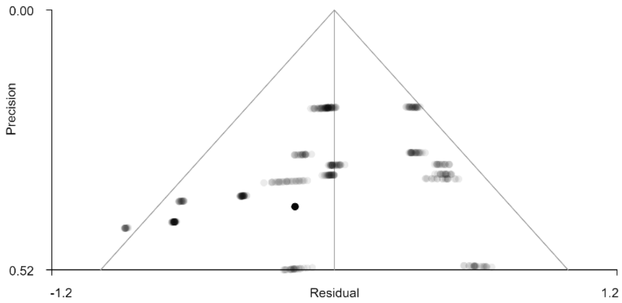

Left inferior occipital gyrus, p=0.368

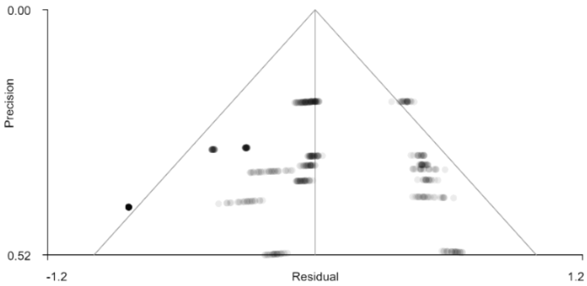

Right inferior parietal gyrus, p=0.735

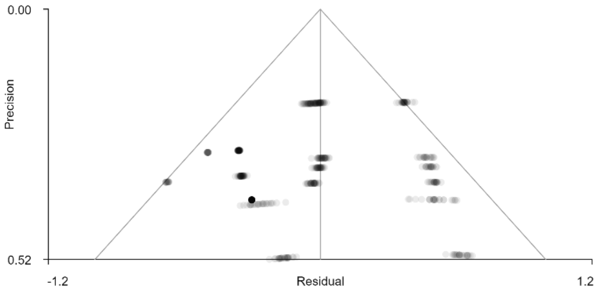

Right middle temporal gyrus, p=0.793

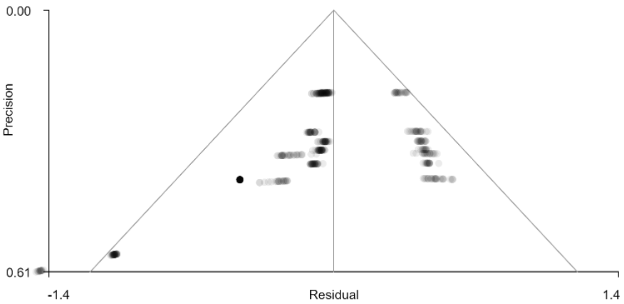

Right cerebellum (hemispheric lobule VIII), p=0.230

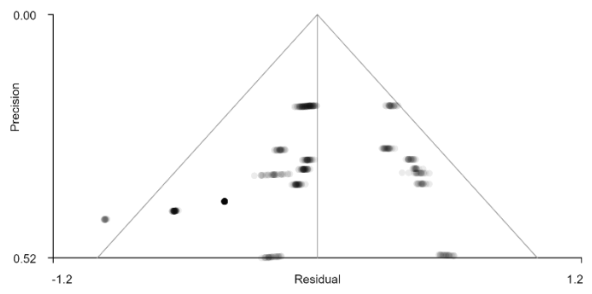

Left median cingulate/paracingulate gyrus, p=0.524

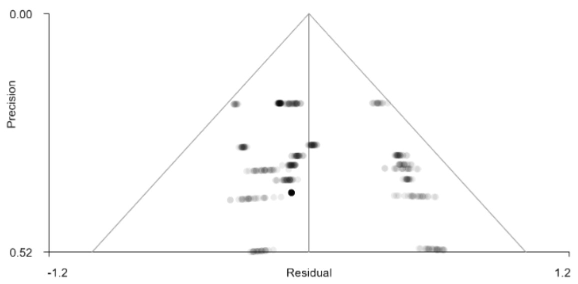

Left inferior temporal gyrus, p=0.611

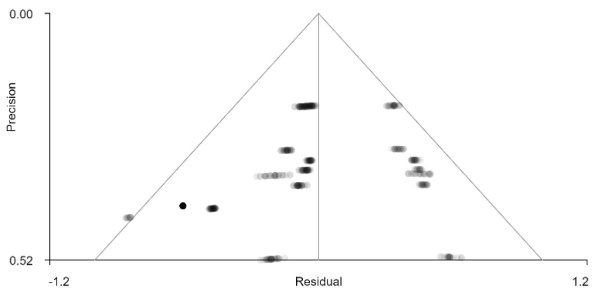

Right supramarginal gyrus, p=0.491

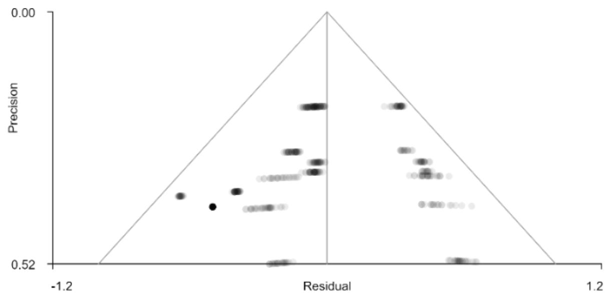

Left postcentral gyrus, p=0.639

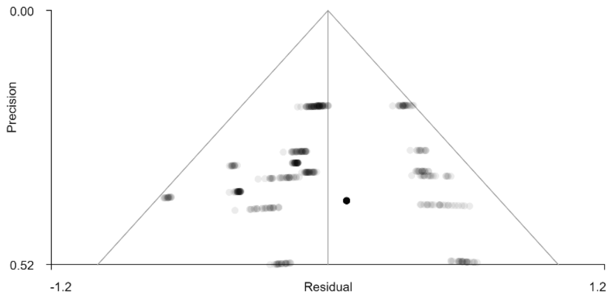

Right lingual gyrus, p=0.842

**eFigure2.** Peak areas with grey matter loss in PDP patients with LEDD (expressed in mg/day) as covariate (uncorrected). This indicated by the blue colour bar on which represent the T threshold of the voxels within the map.

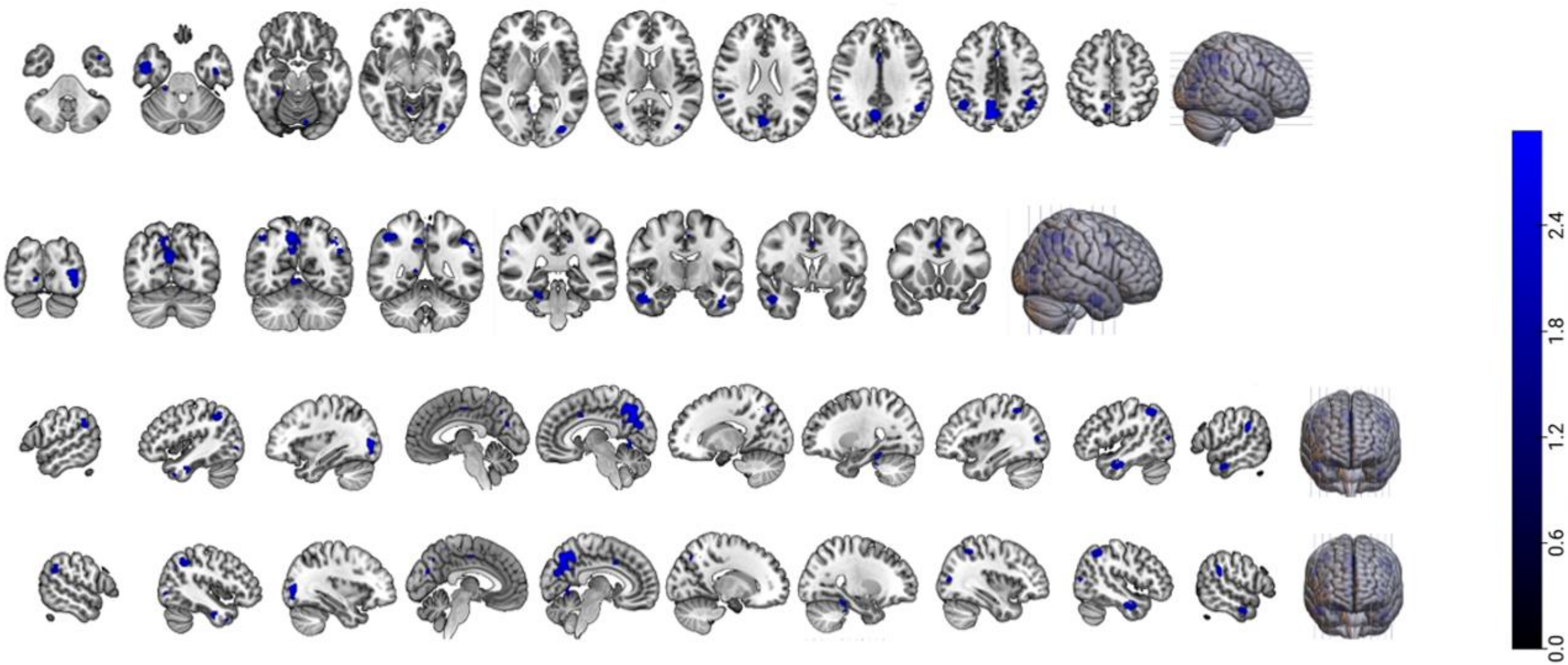

Funnel plots for each peak brain region (with *p* values) with significant grey matter volume loss in PDP patients compared to PDnP patients when PD medications, expressed as LEDD, entered as a covariate. Due to the small number of studies (<10 studies) that were included in the metabias test, all *Z* scores were equal to 0, and *p* values were close to p =1.000.

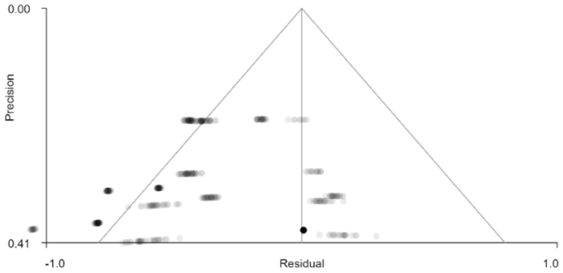

Right precuneus, p = 1.000

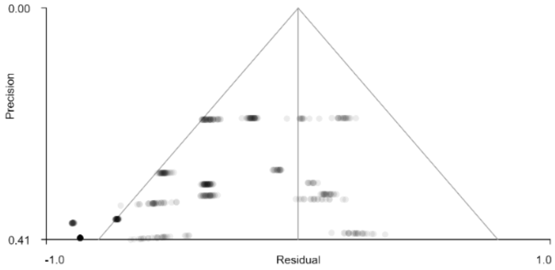

Left angular gyrus, p= 1.000

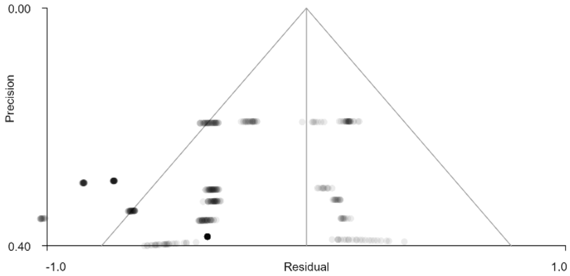

Right inferior temporal gyrus, p=1.000

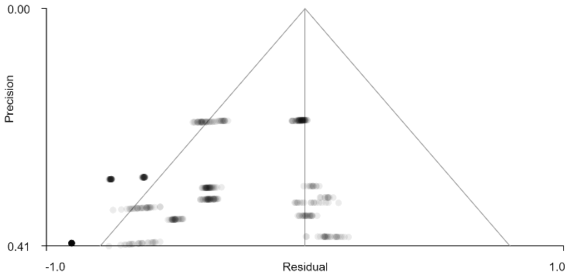

Left middle occipital gyrus, p = 1.000

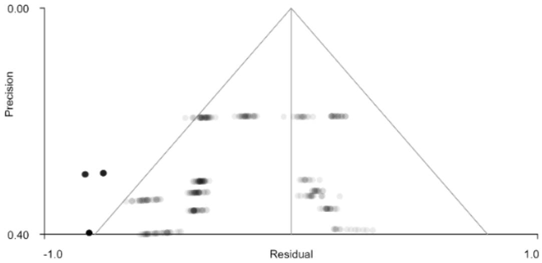

Right inferior parietal gyrus, p=1.000

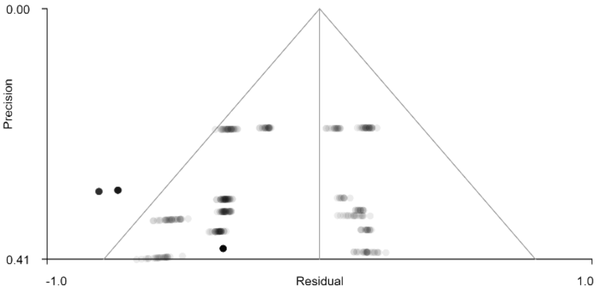

Left medial cingulate/paracingulate gyrus, p=1.000

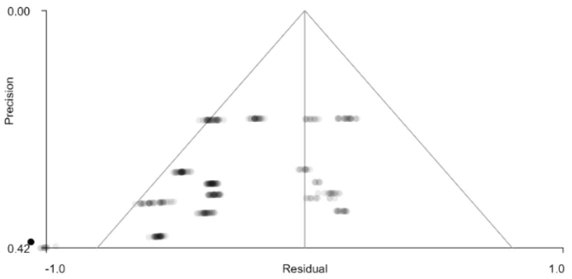

Right supramarginal gyrus, p=1.000

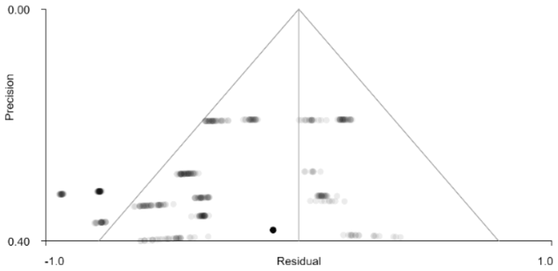

Right lingual gyrus, p=1.000

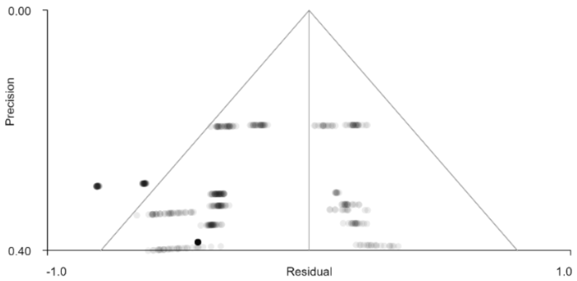

Right middle occipital gyrus, p=1.000

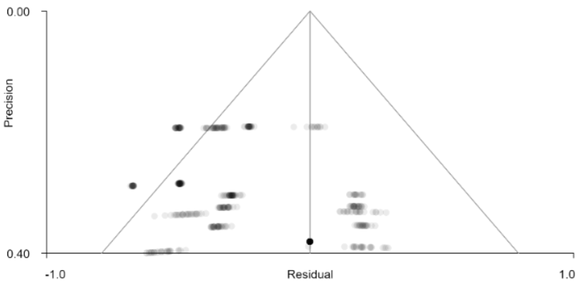

Left inferior temporal gyrus, p=1.000

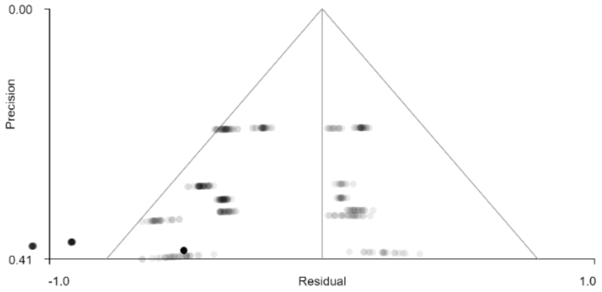

Right fusiform gyrus, p=1.000

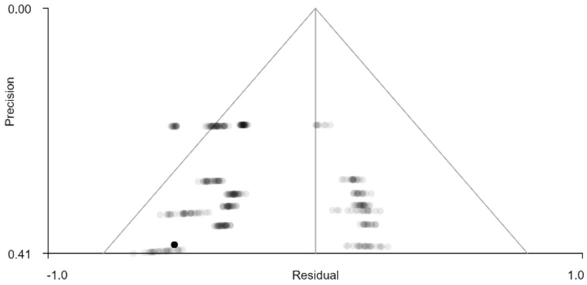

Left temporal pole, p=1.000

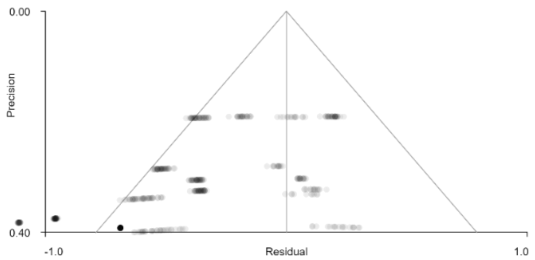

Left inferior parietal gyrus, p=1.000

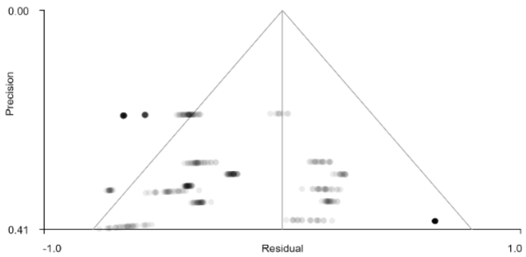

Left cerebellum hemispheric lobule VI, p=1.000

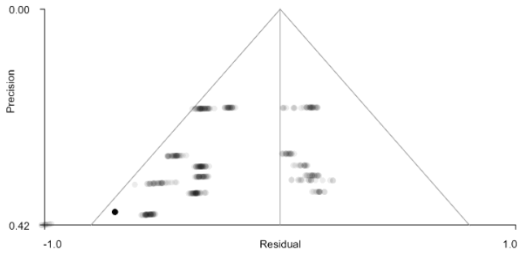

Left supplementary motor area, p=1.000

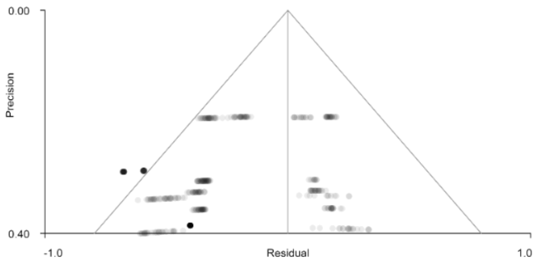

Right calcarine fissure, p=1.000

**eFigure3.** Peak areas with grey matter loss in PDP patients with cognitive scores as covariate (uncorrected). This is indicated by the green colour bar on which represent the T threshold of the voxels within the map.

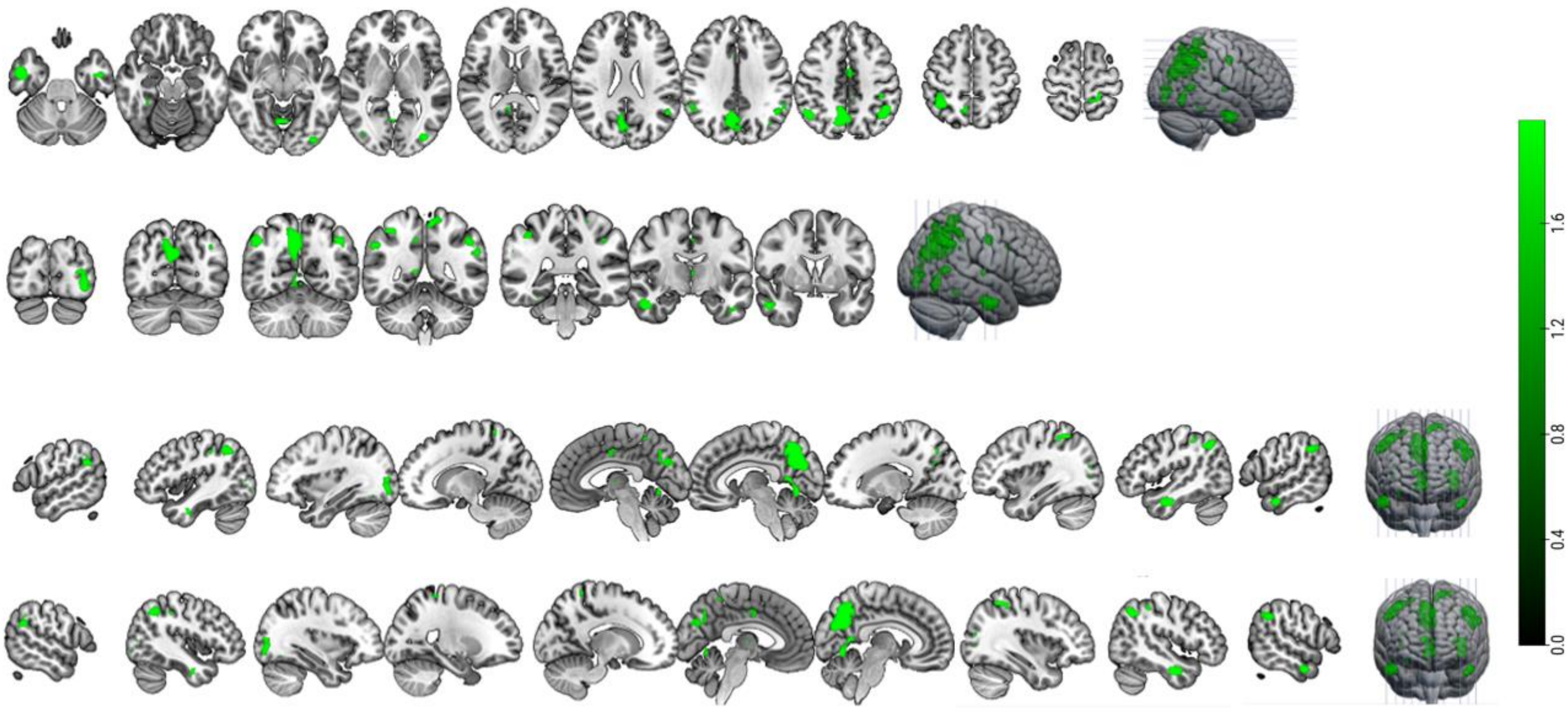

Funnel plots for each peak brain region (with *p* values) with significant grey matter volume loss in PDP patients compared to PDnP patients when cognitive scores entered as a covariate in the analysis. Due to the small number of studies (<10 studies) that were included in the metabias test, all *Z* scores were equal to 0, and *p* values were close to p =1.000.

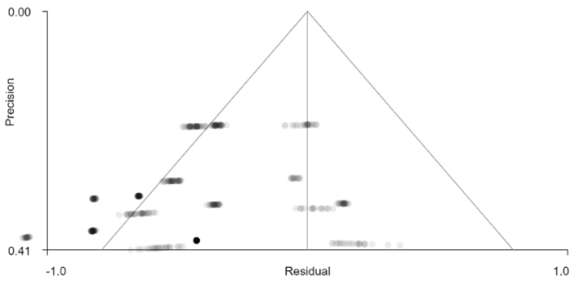

Right precuneus, p = 0.985

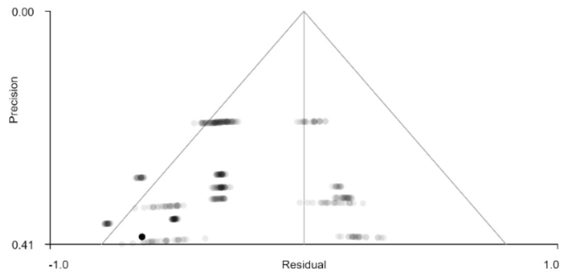

Left inferior parietal gyrus, p = 0.996

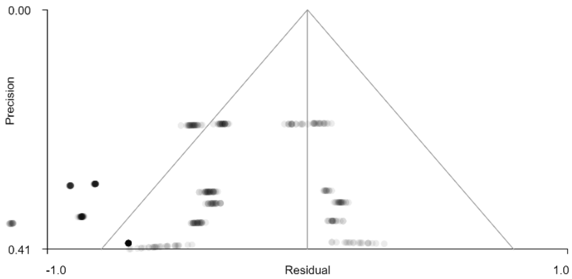

Right inferior temporal gyrus, p = 0.984

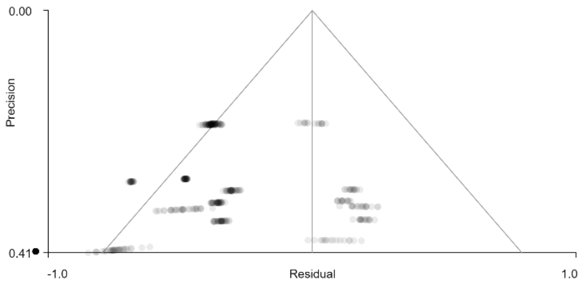

Right inferior parietal gyrus, p = 0.992

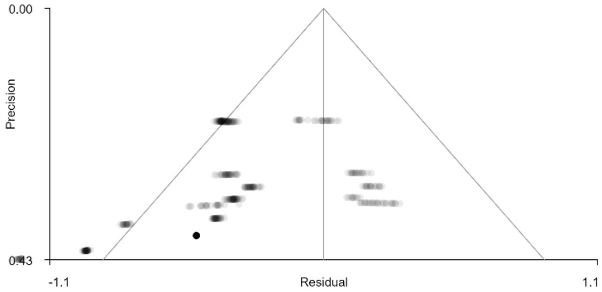

Left inferior occipital gyrus, p = 0.983

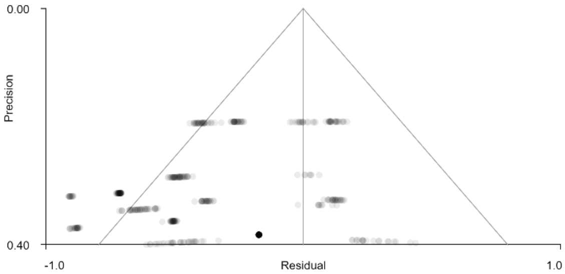

Right lingual gyrus, p =0.978

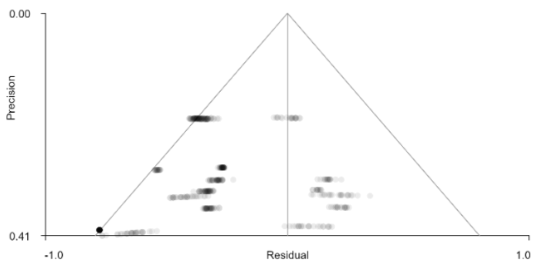

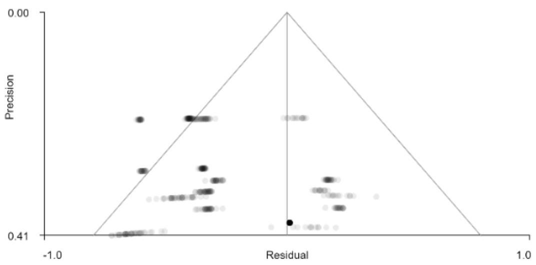

Left postcentral gyrus, p = 0.990

Left inferior temporal gyrus, p = 0.995

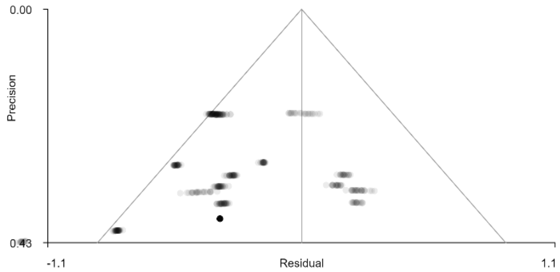

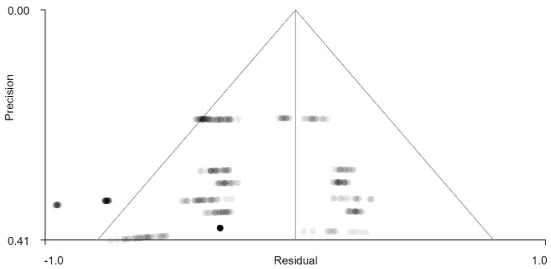

Left median cingulate/paracingulate, p = 0.990

Right superior parietal gyrus, p = 0.985

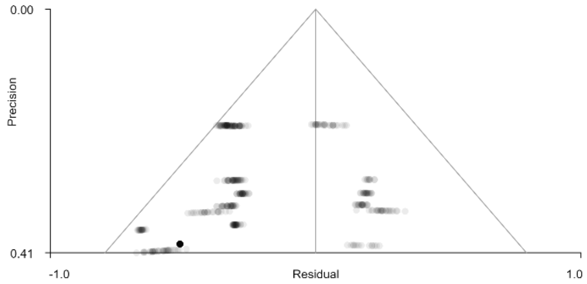

Right fusiform gyrus, p = 0.993

### Supplementary Material 4

**eFigure4.** Scatterplots showing the relationship between D1, 5-HT2a, 5-HT1a receptor gene expressions and Hedges' g effect-size estimates of grey matter volume loss in PDP patients (unadjusted). The regression lines are adjusted for all the predictors in the models.

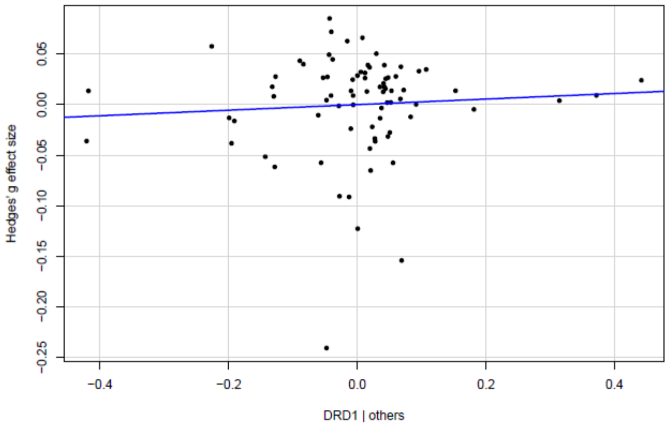

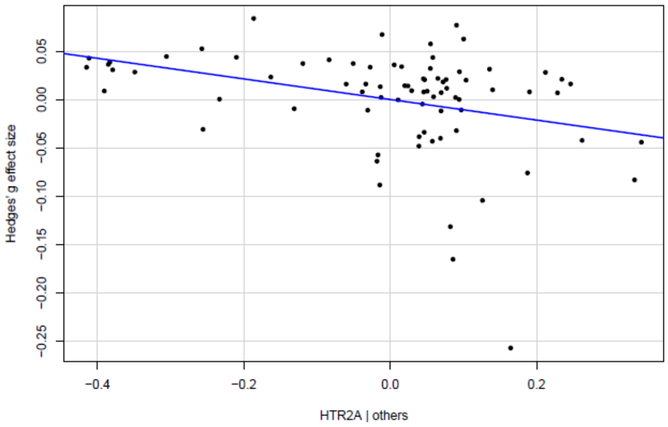

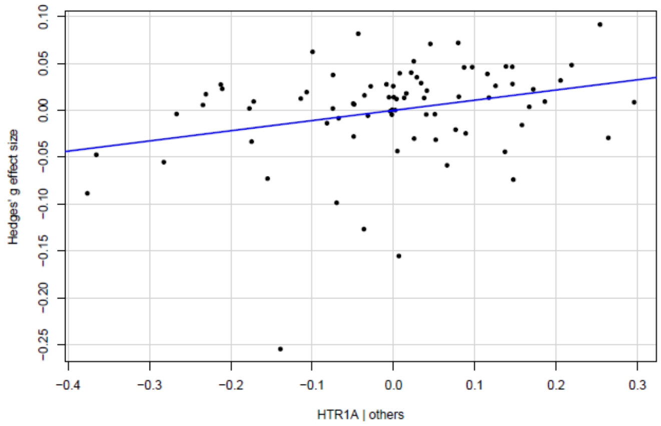

**eFigure5.** Gene expression density of 5-HT2a (A) and 5-HT1a (B) receptors in cortical and subcortical regions, and Hedges' g effect size in cortical and subcortical regions (C) derived from the covariate meta-analysis (unadjusted) results parcellated across the 78 brain regions of the Desikan-Killiany atlas (Desikan et al., 2006).

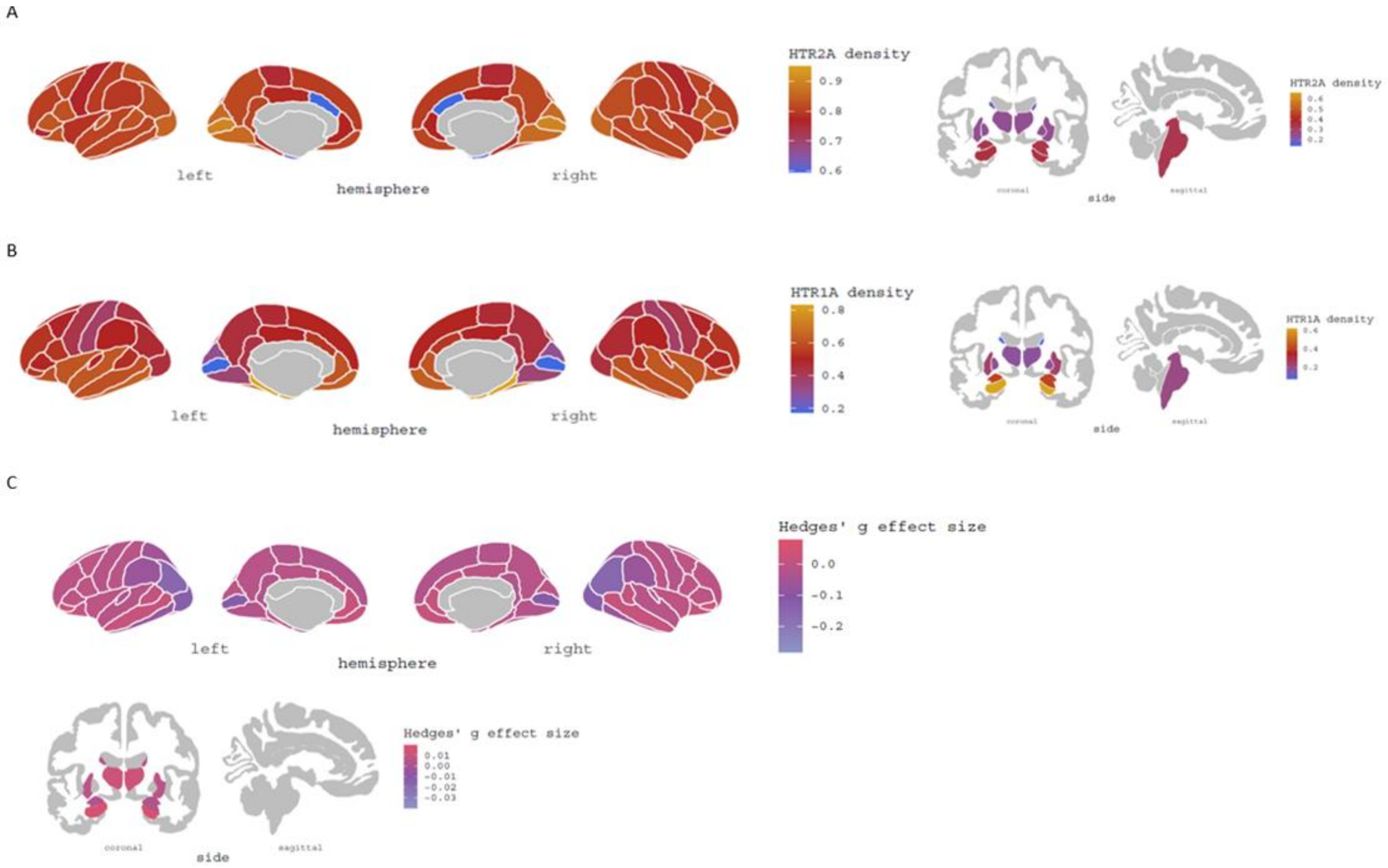
